# Object-centered population coding in CA1 of the hippocampus

**DOI:** 10.1101/2022.07.07.499197

**Authors:** Anne Nagelhus, Sebastian O. Andersson, Soledad Gonzalo Cogno, Edvard I. Moser, May-Britt Moser

## Abstract

Objects and landmarks are crucial for guiding navigation and must be integrated into the cognitive map of space. Studies of object coding in the hippocampus have primarily focused on activity of single cells. Here we record simultaneously from large numbers of hippocampal CA1 neurons to determine how the presence of a salient object in the environment alters single-neuron and neural-population dynamics of the area. Only a small number of cells fired consistently at the object location, or at a fixed distance and direction from it; yet the majority of the cells showed some change in their spatial firing patterns when the object was introduced. At the neural population level, these changes were systematically organized according to the animal’s distance from the object. This organization was widely distributed across the cell sample, suggesting that some features of cognitive maps – including object representation – are best understood as emergent properties of neural populations.

## INTRODUCTION

The mammalian hippocampus and parahippocampal regions are widely believed to construct a cognitive map of physical space^1^. A key element of this map is thought to be place cells in the hippocampus that fire when an animal is in a specific position within an environment^2,3^. Objects and landmarks, which are abundant in real-world environments, are crucial for guiding navigation, and must be part of the cognitive map of space. Consistent with this idea, a long history of work has shown that objects influence spatial firing in the hippocampus in a variety of ways. In place cells, the number, size, position and directionality of firing fields may be changed when a salient object is introduced in the environment^4–9^. Additionally, some studies have reported hippocampal cells with specific firing at the location of the object^5,10^. These ‘object cells’ consistently move their firing fields along with the object when the object is moved and possibly represent the incorporation of objects into the cognitive map of space.

One synapse upstream of the hippocampus, in the medial entorhinal cortex (MEC), we recently reported a specialised cell type that fires not at the location of an object but whenever the animal is at a fixed distance from the object in a given direction (i.e., the cell’s firing location is given by a vector between the animal and the object)^11,12^. In MEC, these ‘object-vector cells’ are present in large numbers and fire independently of the object’s shape, size and identity. A similar set of cells in the hippocampus, called ‘landmark-vector cells’, also appear to fire at fixed distances and directions from objects^10,13,14^. The first report describing these cells in the hippocampus suggested that they only begin to fire at fixed distances and directions from objects after the animal gains experience with the object^13^. However, limited amounts of data made it difficult to characterise the nature of the cells in a systematic manner.

In general, it remains unclear how prevalent object coding is in individual neurons of the hippocampus. Early studies used limited quantitative criteria to identify object cells and landmark-vector cells. Moreover, cells in the hippocampus often show complex responses that, unlike object cells or landmark-vector cells, lack a simple relationship with the object location^7,15^. How is information about objects then represented in the hippocampus? We considered the possibility that information about the presence and location of the object is expressed in the collective dynamics of the hippocampal population in ways that cannot be extracted from the firing of individual neurons^16–18^. To quantify object-related activity in the hippocampus, at the single-cell and neural-population level, we recorded spike activity from almost 1,200 unique principal neurons and up to more than 600 neurons simultaneously in the CA1 area of the hippocampus as rats foraged in an open field environment with and without a discrete 3-dimensional object on the arena floor. First, we show that only a small group of cells fire at the location of the object (object cells) or at a fixed distance and direction from objects (object-vector cells, reminiscient of landmark-vector cells^13^). Among the cells with such firing properties, many showed highly variable tuning across time. Second, we show that while few hippocampal neurons fire at locations defined by the object, many are sensitive to the presence of an object and change their firing patterns in complex and heterogenous ways when the object is introduced, moved, or removed. Finally, at the population level, we show that the complex responses of single cells are brought together into a coherent framework where the population response reliably communicates the animal’s spatial relationship with the object. The population coding is widely distributed across the recorded CA1 cells, suggesting that a neural population perspective may provide a better understanding of how objects are represented in the hippocampus than a perspective based on firing properties of single cells.

## RESULTS

### Object cells and object-vector cells are sparse in the hippocampus

To investigate single-cell and population coding of objects in the hippocampus, we implanted tetrodes or Neuropixels probes in the CA1 of rats (Supplementary Fig. 1; 7 tetrode rats and 2 Neuropixels rats) and recorded activity daily for four trials as the rats foraged in a 150 cm x 150 cm open field arena (Fig. 1a, left). Each session began with an empty box trial (No Object). Then, on the second trial (Object), a multi-coloured Duplo tower 52 cm × 10 cm × 10 cm (height × width × length) was placed in the arena. On the third trial (Object Moved), the object was moved to a new, semi-random location, before it was taken out from the box for the last trial (No Object’).

**Figure 1.**
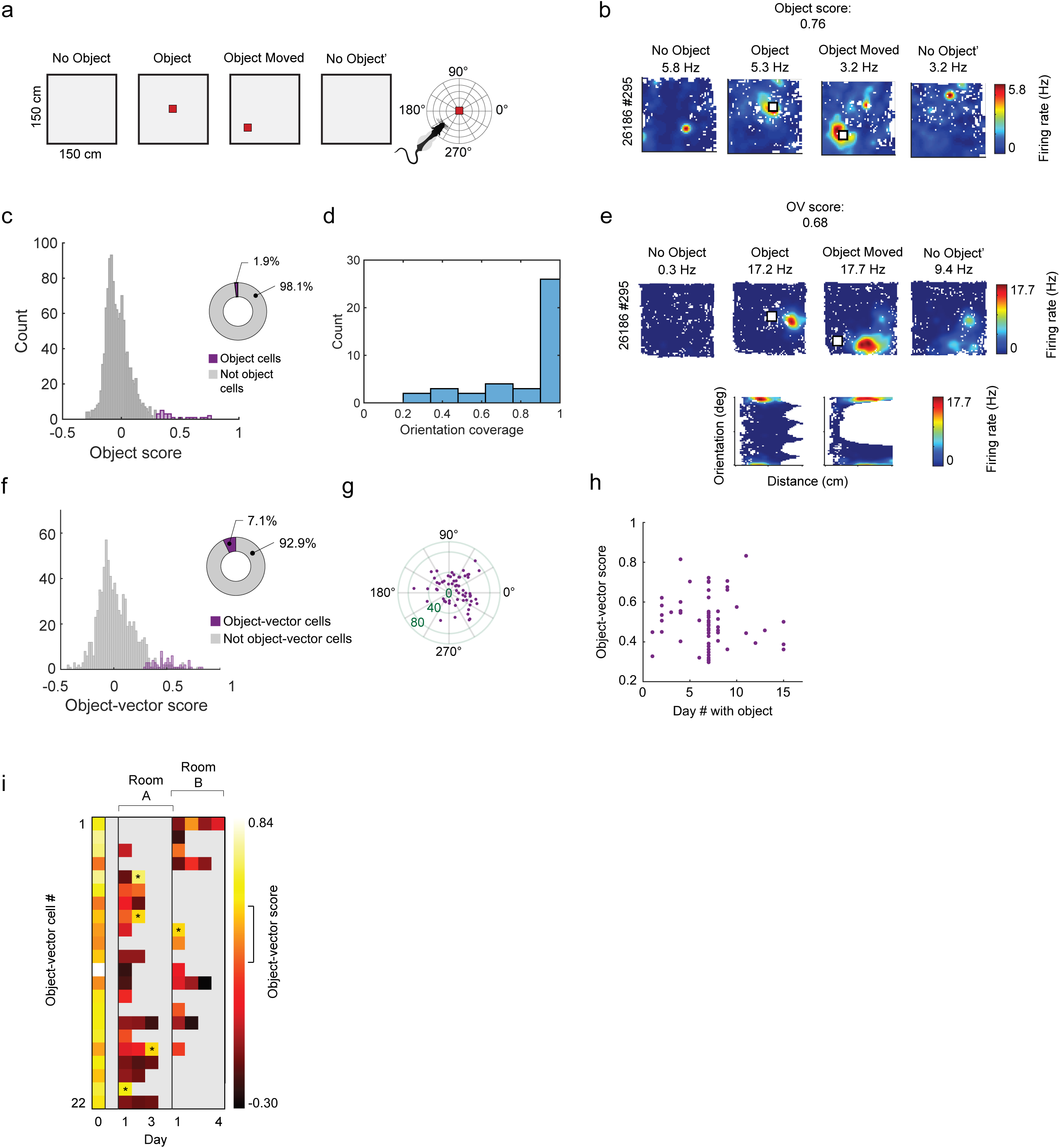
A small number of object cells and object-vector cells are present in CA1. **a,** Left, schematic of experimental protocol (bird’s-eye view). Each recording session consisted of four consecutive trials where rats foraged freely in the same 150 cm x 150 cm open field arena. In the two middle trials (Object and Object Moved), a Duplo tower (height x width x length, 52 cm x 10 cm x 10 cm) was placed into the environment (indicated by the red squares). Right, the distance and orientation from the rat to the object were defined in an allocentric frame with reference to the centre of the object, with east at 0°. Rat cartoon is from http://doi.org/10.5281/zenodo.3926343. **b,** Color-coded firing rate maps from all trials from example object cell (color scale to the right). Object location in the rate map is indicated by the white square. Note that cell fires at the location of the object in both the ’Object’ and ‘Object Moved’ trials. Rat ID and cell ID are indicated at the left and peak firing rate at the top. **c,** Left, distribution of object scores (*n* = 1189 unique CA1 cells). Purple bars, cells classified as object cells; grey bars, cells not classified as object cells. Right, pie chart depicting the percentage of object cells (*n* = 23 cells, 1.9%). **d,** Orientation coverage of object cells (n = 46, 23 cells × two trials). The orientation coverage is expressed as the fraction of the entire azimuthal range around the object (0-360°) covered by the firing fields of object cells. Object cells mostly fired on all sides of the object. **e,** Colour-coded firing rate maps from all trials (top) and vector maps from object trials (bottom) from an example object-vector cell. Rate map conventions are as in Fig. 1b. Vector maps show firing rate as a function of distance (x-axis) and allocentric orientation (y-axis) to the object. Locations not sampled are in white. Note that the cell fires in a fixed distance and direction from the object. **f,** Same as for panel c but for object-vector cells (n = 60 cells from a total of 844 CA1 cells were classified as object-vector cells, 7.1%). **g,** Polar scatter plot showing orientation (polar axis, in degrees) and distance (radial axis, in cm, indicated in green) of object-vector fields (one dot per cell). Orientation and distance are calculated from the centre-of-mass of the fields with respect to the object location. **h.** Object-vector score as a function of the number of days with exposure to the object (Day #) for all recorded object-vector cells (*n* = 60). Each dot indicates a cell. **i,** Color-coded matrix showing object-vector scores of 22 object-vector cells that were recorded on more than one day. Left column, object-vector scores for the 22 cells on the first day they were recorded (‘Day 0’). Middle and right column, object-vector scores for the same cells on additional days in the same room (Room A; up to 3 days) or in a different room (Room B; up to 4 days). Only 5 of the 22 cells satisfied criteria for object-vector cells on more than one day (cells indicated by asterisks). Grey squares indicate that the cell was not recorded on the specific day. Bar, range of object-vector scores (lighter colours, high object-vector score; darker colours, low object-vector score). The range of 99^th^ percentiles of the shuffled distributions for object-vector score from all cells is indicated by the line to the right of the bar (range: 0.27-0.48).

To identify object cells, which fire at the location of the object^5,10^, we used a template-matching procedure. First, we placed a Gaussian firing field at the location of the object as the template for an ideal ‘object cell’. Second, we calculated the Pearson correlation between the firing rate map of the cell and the template. This was done both for the Object and Object Moved trials (using separate templates for each trial). We then took the minimum of the two Pearson correlations as an ’object score’, which will be high for cells firing at the object’s location in both trials (Fig. 1b). To calculate how many object cells were present in our dataset, we used a shuffling procedure where, for each iteration, we randomly chose the templates’ locations and re-calculated the score. Cells with object scores above the 99th percentile of the shuffling distribution were considered object cells. Only 1.93% of the CA1 cells classified as object cells (*n* = 23/1189 cells). Although low, this is significantly above a chance level of 1% (*p* = 0.0026, binomial test) (Fig. 1c) (Supplementary Fig. 2). Object cells mostly had firing fields covering the entire range of orientations around the object (Fig. 1d) and did not display a consistent orientation preference relative to the centre of the object across trials (change in orientation of centres-of-mass of firing fields between Object and Object Moved, mean ± SD: 74.3 ± 49.4°). This confirms the existence of object cells in CA1 as previously described^5,10^ but suggests that they exist only in small numbers.

We next asked how many hippocampal cells fire at fixed distances and directions from the object (i.e. show vectorial firing relative to the object), rather than at the object’s location. Previously, such object-vector cells have been shown to be abundant in MEC^11,12^. However, in the hippocampus (where they have been referred to as landmark-vector cells^13^) they have been reported only in small numbers^5,10^. As a first step to identify object-vector cells in the larger sample of the present data, we expressed the activity of cells in ‘vector maps’, following procedures established for MEC cells^11,12^, where firing rate is shown as a function of allocentric distance and orientation to the object (Fig. 1a, right; Fig. 1e, bottom). We then used an object-vector score to identify cells that fire at a fixed distance and direction from the object^11,12^. The object-vector score is the Pearson correlation between the vector maps from the Object and Object Moved trials (Fig. 1e, bottom), which is high when the cell’s firing is determined by the rat’s position relative to the object. We created shuffled data by randomly shifting the spiketrains in time and considered cells to be object-vector cells if they had object-vector scores higher than the 99 percentile of the shuffled distribution (and if they passed additional criteria used previously to identify object-vector cells^11,12^, see Method for details). In CA1, 7.1% of all cells were classified as object-vector cells (*n* = 60/844 cells; significantly above a chance level of 1%, *p* = 2.32×10^−31^, binomial test) (Fig. 1f, Supplementary Fig. 3). We confirmed with an independent region-of-interest method that these cells indeed fired at fixed distances and directions from objects (Supplementary Fig. 3b, c). As in MEC^11^, firing fields of the cells covered a range of distances from the object (15.2-80.7 cm; mean ± SD: 37.7 ± 16.0 cm) and the entire azimuthal range (Fig. 1g). Taken together, we confirm both the existence of object cells and object-vector cells in CA1, but only in small numbers.

The first report on landmark-vector cells (here called object-vector cells) suggested that the cells’ response became stable only after the animal had gained sufficient experience with the object^13^. Could slow acquisition of vectorial firing account for the small numbers of object-vector cells in our data? This was not the case. We found object-vector cells both when the animals had no experience with the object and when they were very familiar with the object (Fig. 1h) and observed no significant effect of day of exposure to the object on either the object-vector score (Fig. 1h) (linear regression, *R^2^* = 0.01, *p* = 0.50) or the percentage of object-vector cells (Supplementary Fig. 3d) (Pearson correlation, *r* = −0.009, d. f. = 14, *p* = 0.97). This suggests that object-vector responses in CA1 can be present already from the first trial and that the low number of object-vector cells is not explained merely by lack of experience with the object. The sample of object cells recorded across different days was too small to perform the same analysis as for object vector cells, but we observed that object cells could also be present from the first day of exposure to the object.

We next asked if the object cells’ responses were stable across time and environments. Within the total sample of 60 object-vector cells, we were able to record a subset of 22 cells across more than one day (in two different rooms, Room A and Room B). In many cells, the object-vector score appeared unstable, decaying from high values on the first day they were identified as object-vector cells (Fig. 1i, leftmost column) to lower values on later days (Fig. 1i, all columns for Room A and B). Only 5 cells had significant object-vector scores on one or more subsequent days (Fig. 1i, cells indicated by asterisks). Despite this instability across time, object-vector cells were more stable than expected by chance, showing a higher probability of passing the criteria for object-vector cells on later days compared to other cells (5/50 cases; probability = 0.1 vs. 2/219 cases; probability = 0.0091; *p* = 9.40 x 10^−5^, binomial test). It was not possible to determine the stability of object cells in a similar way due to low cell numbers. During the course of a single trial, the responses of object cells were only moderately stable (Pearson correlation between rate maps from 1st and 2nd halves of trial, mean ± SEM: 0.50 ± 0.06 for ‘Object’ trial; 0.48 ± 0.05 for ‘Object Moved’ trial), consistent with the possibility that they too display instability across time.

Finally, we showed that object-related activity is also present in the CA3^13,19^. Recording from the CA3 and hilus in 4 rats (Supplementary Fig. 1), we found that 2.0% of cells were object cells and 3.7% of cells were object-vector cells (Supplementary Fig. 4a-d) Altogether, this indicates that object cells and object-vector cells are sparse in CA1, but possibly even more so in CA3, consistent with previous reports^14^.

### The majority of hippocampal cells change spatial firing patterns in the presence of objects

The number of object cells and object-vector cells was low in our data, but this does not necessarily mean that most CA1 cells do not respond to the presence of an object. We found numerous examples of CA1 cells changing firing patterns at all stages of our experimental protocol (Fig. 2a), although these changes differed from those expected of object cells and object-vector cells. To probe the changes quantitatively, we asked how much the spatial firing patterns and firing rates changed as a consequence of object introduction (No Object vs. Object), object movement (Object vs. Object Moved), and object removal (Object Moved vs. No Object’). For each cell (1189 principal neurons in CA1 from 9 rats), we calculated the Pearson correlation between the rate maps to assess changes in the spatial firing patterns. To assess changes in firing rate, we used a measure of rate overlap^20,21^, defined as the mean firing rate in the less active trial divided by the mean firing rate in the more active trial.

**Figure 2.**
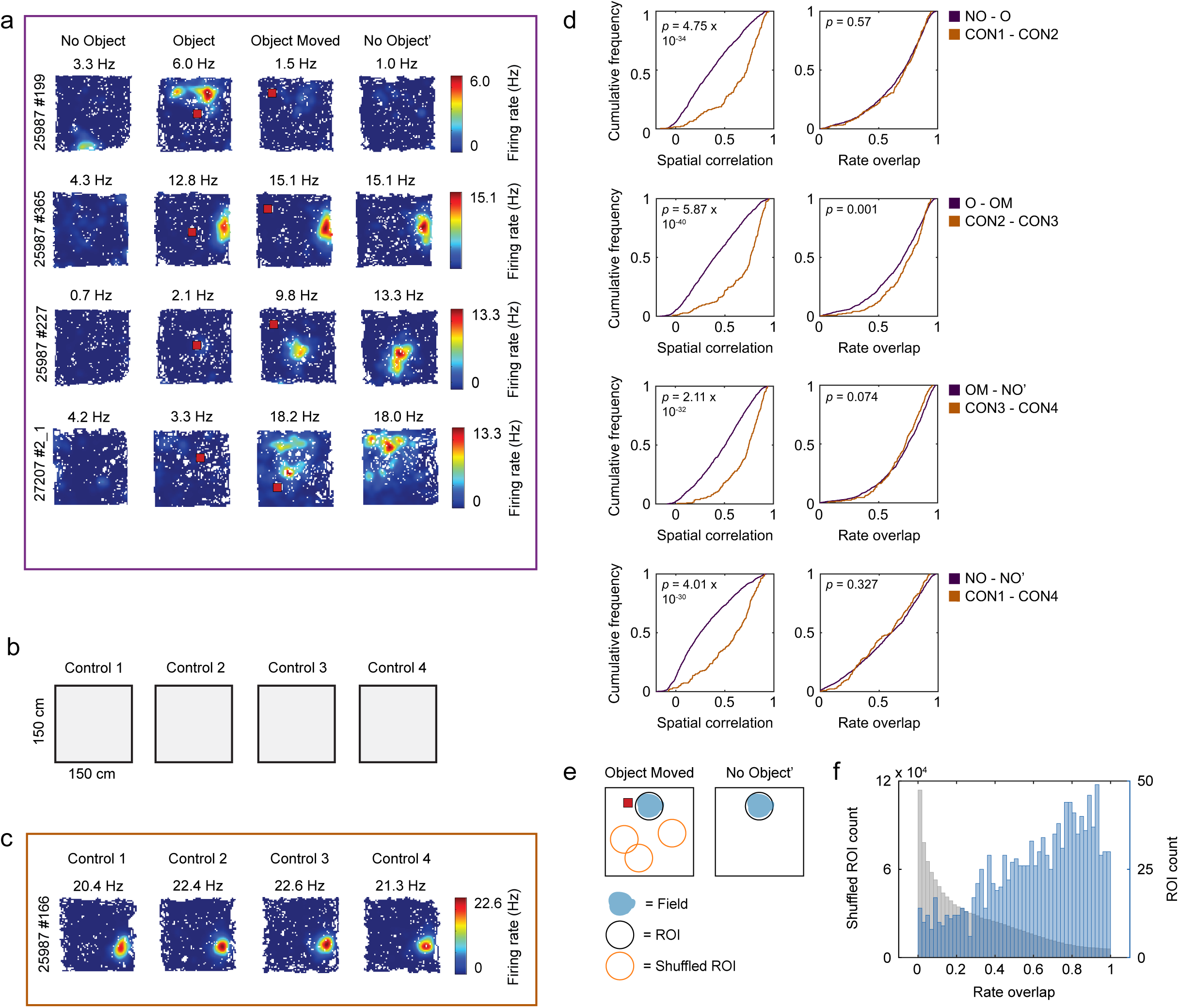
CA1 cells change spatial firing patterns in response to objects. **a.** Cells responded by changing their spatial firing patterns when an object was introduced, moved, or taken out of the box. Colour-coded firing rate maps of 4 example cells recorded in CA1 (experimental protocol as in Fig. 1a). The first cell changes its spatial firing pattern only upon introduction of the object (top row). The second cell also changes its spatial firing pattern when the object is introduced but it preserves the firing pattern during the next trials, leaving a trace of the original firing field (second row). The third cell changes its firing pattern when the object is moved and maintains a trace on the last trial (third row). The fourth cell changes its firing pattern when the object is moved, but the spatial firing pattern is more diffuse than in the other cells (last row). Object locations are shown as red squares. Rat ID and cell ID are indicated to the left of the firing rate maps, and peak firing rate in each trial is indicated at the top of each rate map. Bars, colour-coded firing rate. **b.** Control experiments used to compare object-induced changes to changes that occur due to passage of time. Schematic of experimental protocol (bird’s-eye view). Each recording session consisted of four consecutive trials where rats foraged freely in the same 150 cm x 150 cm empty open field arena. **c.** Colour-coded firing rate map from an example CA1 cell in the control experiment. The cell does not change its activity throughout the session. Symbols as in panel a. **d.** Cumulative normalized frequency distributions showing spatial correlation (left column) and rate overlap (right column) between pairs of trials. Each row shows a given pair of trials in sessions with Objects (as in Fig 1a; dark purple lines; *n* = 1189 principal cells recorded from 9 rats) and the corresponding pair of control trials (as in Fig. 2b; brown lines; *n* = 219 principal cells recorded from 3 rats). The spatial correlation is defined as the Pearson correlation between the firing rate maps of the two indicated trials (e.g., NO vs. O), and the rate overlap is calculated by dividing the mean firing rate in the less active of these trials by the mean firing rate in the more active trial. NO = No Object; O = Object; OM = Object Moved; NO’ = No Object’; CON 1-4 = Control 1-4. **e.** Schematic of procedure to quantify object trace responses. First, we identified cells where the spatial firing pattern changed between the first and last trials (No Object and No Object’; *n* = 919 cells). In these cells, we identified firing fields in No Object’ as a circular region-of-interest (ROI; ‘Field’, blue). We then transferred the ROI to the same location in the Object Moved trial and calculated the rate overlap between the ROIs. To create shuffled data, we moved the ROI to random locations in the Object Moved trial (‘Shuffled ROI’, orange) and calculated the rate overlap between the actual ROI in No Object’ and the Shuffled ROI in Object Moved. This shuffling procedure was repeated 1000 times. If the cell had more than one field, we repeated the analysis for each field. **f**. Rate overlap values from shuffled data (left y-axis, gray) and real data (right y-axis, blue). Distribution for real data is shifted to the right compared to shuffled data, showing that in many CA1 cells firing fields persist after the object is removed.

First, we assessed the changes in spatial firing patterns and firing rates after introducing the object (No Object – Object trials). Since place cells change activity merely due to passage of time^22–24^, we compared the change elicited by the presence of the object with a condition where the rats ran in an open environment without an object present (Fig. 2b, c; Control 1 – 4; 219 unique principal cells from 3 rats). Introducing the object led to larger changes in spatial firing patterns (i.e., smaller spatial correlations) than in the corresponding control trials (Fig. 2d, top left) (NO – O vs. CON1 – CON2: *z =* 12.17, *p* = 4.75 x 10^−34^, Wilcoxon rank sum test), while the degree of rate change was the same (Fig. 2d, top right) (NO – O vs. CON1 – CON2: *z =* 0.57, *p* = 0.57, Wilcoxon rank sum test). Using a shuffling procedure (see Method for details), we estimated that 67.7% of the cells (*n* = 803/1189 cells) express some type of change between No Object and Object – that is, the majority of cells respond to introduction of the object.

Second, we examined changes between the Object and Object Moved trials, i.e., when the object was moved from one location in the environment to another. Here too, CA1 cells responded with large changes in spatial firing patterns compared to control trials (Fig. 2d, second row, left) (O – OM vs. CON2 – CON3: z = 13.23, *p* = 5.87 x 10^−40^), whereas the amount of rate change was again small (O – OM vs. CON2 – CON3: Fig. 2d, second row, right) (*z =* 3.43, *p* = 0.001, Wilcoxon rank sum tests). In total, we estimated that 69.9% of the cells changed their firing when the object was moved (*n* = 829/1,186 cells). Finally, when we removed the object, the cells changed their spatial firing patterns, but not firing rates, compared to control trials (Fig. 2d, third row) (OM – NO’ vs. CON3 – CON4; spatial correlation: *z* = 11.85, *p* = 2.11 x 10^−32^; rate overlap: *z* = 1.79, *p* = 0.074, Wilcoxon rank sum tests). We observed similar results in CA3, although the CA3 cells were less sensitive to the presence of the object than CA1 cells (Supplementary Fig. 4g).

At the same time, many cells preserved firing fields between the Object Moved and No Object’ trials (Fig. 2a, middle rows), consistent with the finding that place cells can fire at past locations of an object^20,21,25^. In fact, changes between No Object and No Object’ were greater than between the Object Moved and No Object’ trials, both for spatial and rate changes (Supplementary Fig. 5a) (spatial correlation: *z* = 14.32, *p* = 1.64 x 10^−46^; rate overlap: *z* = 15.35, *p* = 3.40 x 10^−53^, Wilcoxon signed-rank tests). The changes between No Object and No Object’ were also greater than in control trials with no objects (Fig. 2d, bottom row) (NO – NO’ vs. CON1 – CON4: *z* = 11.85, *p* = 2.11 x 10^−32^, Wilcoxon rank sum test), confirming the presence of hysteresis effects in CA1 place cells.

To directly quantify the presence of object traces, we used an ROI-based approach (Fig. 2e, f). For each cell, we first identified firing fields in the No Object’ trial as ROIs (Fig. 2e, right, “Field” in blue). We then transferred each ROI to the same location in the Object Moved trial and calculated the rate overlap between the ROIs (Fig. 2e, left). To create shuffled data for each cell, we moved the ROI to random locations in the box (Fig. 2e, left, “Shuffled ROI” in orange) and calculated the rate overlap with the original ROI in the No Object’ trial. If the cell had more than one field in the No Object’ trial, we repeated the analysis for all fields (*n* = 1,252 fields in *n* = 919 cells). Using the ROI method, we found that 23.7% of CA1 cells had trace responses, with larger values of rate overlap than the 99^th^ percentile of shuffled data (Fig. 2f and Supplementary Fig. 5) (*n* = 218/919 cells in total, significantly above chance level of 1%, *p* < 5.73×10^−141^, binomial test). Taken together, we find that objects strongly influence the firing of CA1 cells in a variety of ways, both when the object is present and after it has been removed.

### Decoding the presence of an object from neural population activity

So far we have shown that a large proportion of cells change their firing pattern when an object is introduced, moved or removed in the environment, indicating that hippocampal responses are far from agnostic to the object’s presence. However, hippocampal responses appear different from those in MEC, where a single functional cell type, object-vector cells^11^, exists in large numbers and might play a dominant role in object representation. By contrast, in CA1 of the hippocampus, we find that cells show heterogenous responses (Fig. 1–2), much like recently described in the dentate gyrus^15^. This made us wonder if the heterogeneous responses of single cells are brought together and organised in a meaningful way when viewed as a collective whole. That is, do we obtain a different understanding of the hippocampal code for object representation when analysing large populations of cells recorded simultaneously?

We first reasoned that if hippocampal population activity contains object-related information, we would be able to use population activity to decode whether an object is present in the environment. For all decoding analyses, we used recordings from 15 tetrode sessions with 25-39 cells in each recording and 2 Neuropixels sessions with 620 and 135 cells, respectively, unless otherwise indicated. Building upon an existing Bayesian decoding algorithm^26^, we developed an approach where we feed data from the No Object trial and the Object trial into a decoder, which classifies testing data into either of these two types of trial (Fig. 3a). Specifically, each sample of testing data was a vector of spike counts from all simultaneously recorded hippocampal cells in a 1 second time window. The decoding accuracy was above chance (50%) in all experimental sessions (Fig. 3b), showing that we can infer the presence of the object using population activity. The decoder’s performance increased approximately logarithmically with the bin size of the testing data vector (Fig. 3c, d) and the success of decoding increased with the number of cells included (Fig. 3e) (decoding accuracy with 10 cells: 58.3% ± 0.81%; decoding accuracy with 100 cells: 77.5% ± 1.07%; decoding accuracy with 500 cells: 85.2% ± 1.50%; mean ± SEM; quantification based on Neuropixels dataset with N = 620 cells in total). We chose 1 second bins for the main analyses because performance saturated around this level (Fig. 3c).

**Figure 3.**
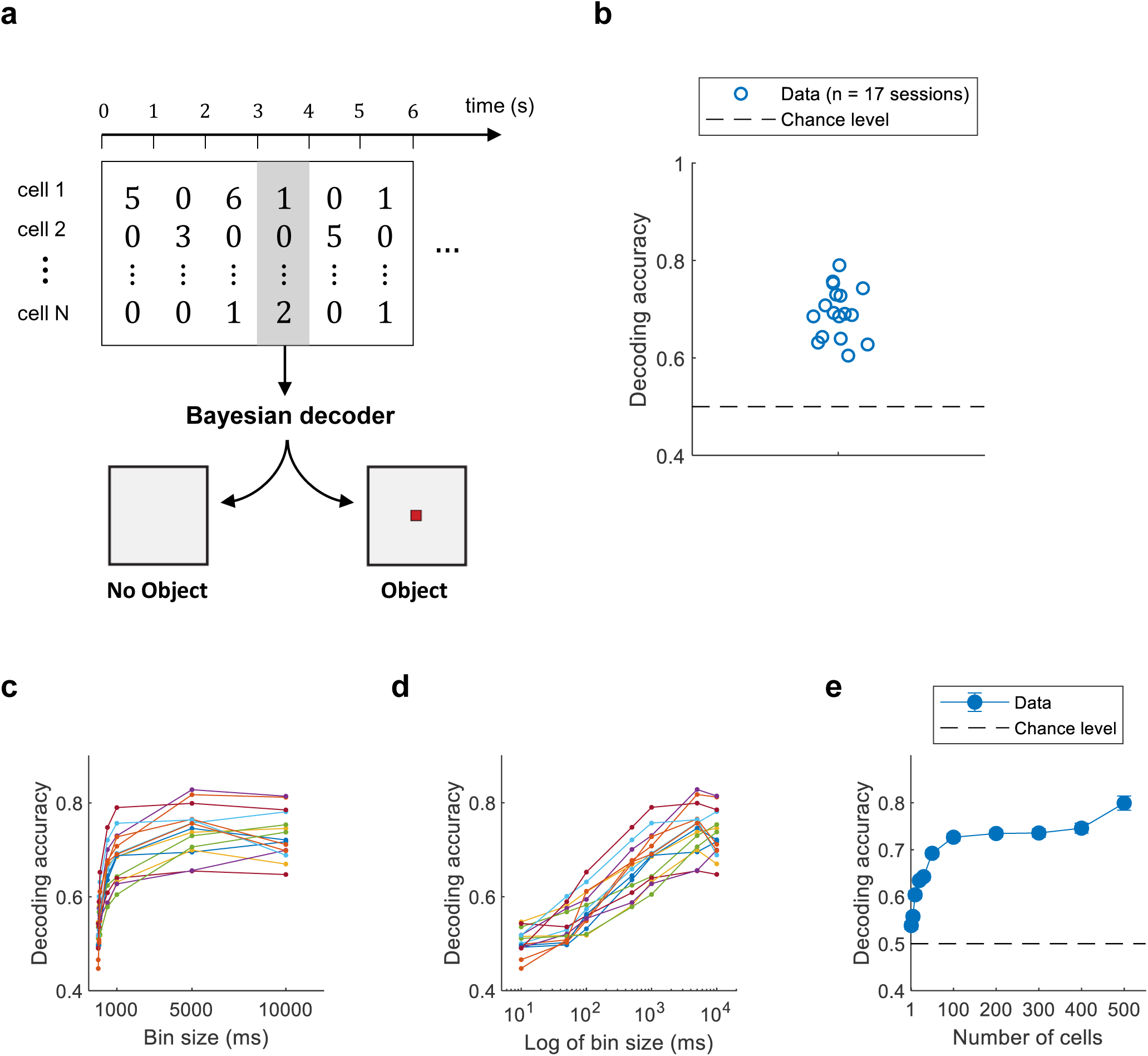
Decoding the presence of an object in the animal’s environment. **a,** Schematic of the conceptual framework. We take out a sample of population data (highlighted in gray), consisting of spike counts of simultaneously recorded hippocampal cells in a 1 second time window, and ask whether a Bayesian decoder can infer whether this 1 second sample of data occurred in the absence or presence of an object (left and right, respectively). **b,** Decoding accuracy in all sessions (n = 17). The decoding accuracy is given by the fraction of data samples that the decoder classified correctly. Stippled line indicates chance level (0.5). **c,** Decoding accuracy as a function of bin size (ms). The bin size varied between 10, 50, 100, 500, 1000, 5000 and 10000 ms. Performance saturates around 1000 ms, which motivated our choice of 1 second bins for the main analysis. Each color shows a different experimental session. **d,** Same data as in b but with x-axis rescaled to show linear increase in performance as a function of logarithm of bin size. **e,** Decoding accuracy as a function of number of cells used to perform the decoding. Data were generated by taking a random subset of n cells from a total population of 620 simultaneously recorded cells (Neuropixels data, rat 27207) and then running the decoder. For each specific n, we repeated the random subsampling 100 times. Data points show means; error bars show SEM.

Having established as a proof-of-concept that we can use population activity to decode the presence of the object, we next asked if decoding accuracy differed as a function of the animal’s spatial relationship with the object. We reasoned that if decoding accuracy depends on where the animal is with respect to the object, this might provide important clues about how objects in the proximal environment affect neural population activity. We divided the environment into 5 distance bins, consisting of concentric rings radiating out from the object (Fig. 4a). By feeding data from each distance bin into the decoder, we found that decoding performance decreased gradually with increasing distance from the object (Fig. 4b) (*H* = 44.559, *p* = 4.91×10^−9^, Kruskal-Wallis test, n = 17 experimental sessions; significantly different decoding accuracies for distance bins 1 vs. 2; 2 vs. 3; and 4 vs. 5, *W* ≥ 111, *p* ≤ 0.0049, Wilcoxon signed-rank tests). Such a decrease was also seen in the individual experiments (Fig. 4c). In the main analyses, we optimised distance bins for each experiment so that we had an equal number of samples in each bin, avoiding potential biases where the animal might have spent more time in some bins. We found the same decrease in decoding accuracy using fixed distance bins rather than optimised bins (Supplementary Fig. 6a and b) (fixed bins of 0-20 cm; 20-40 cm; 40-60 cm; 60-80cm; 80-100 cm).

**Figure 4.**
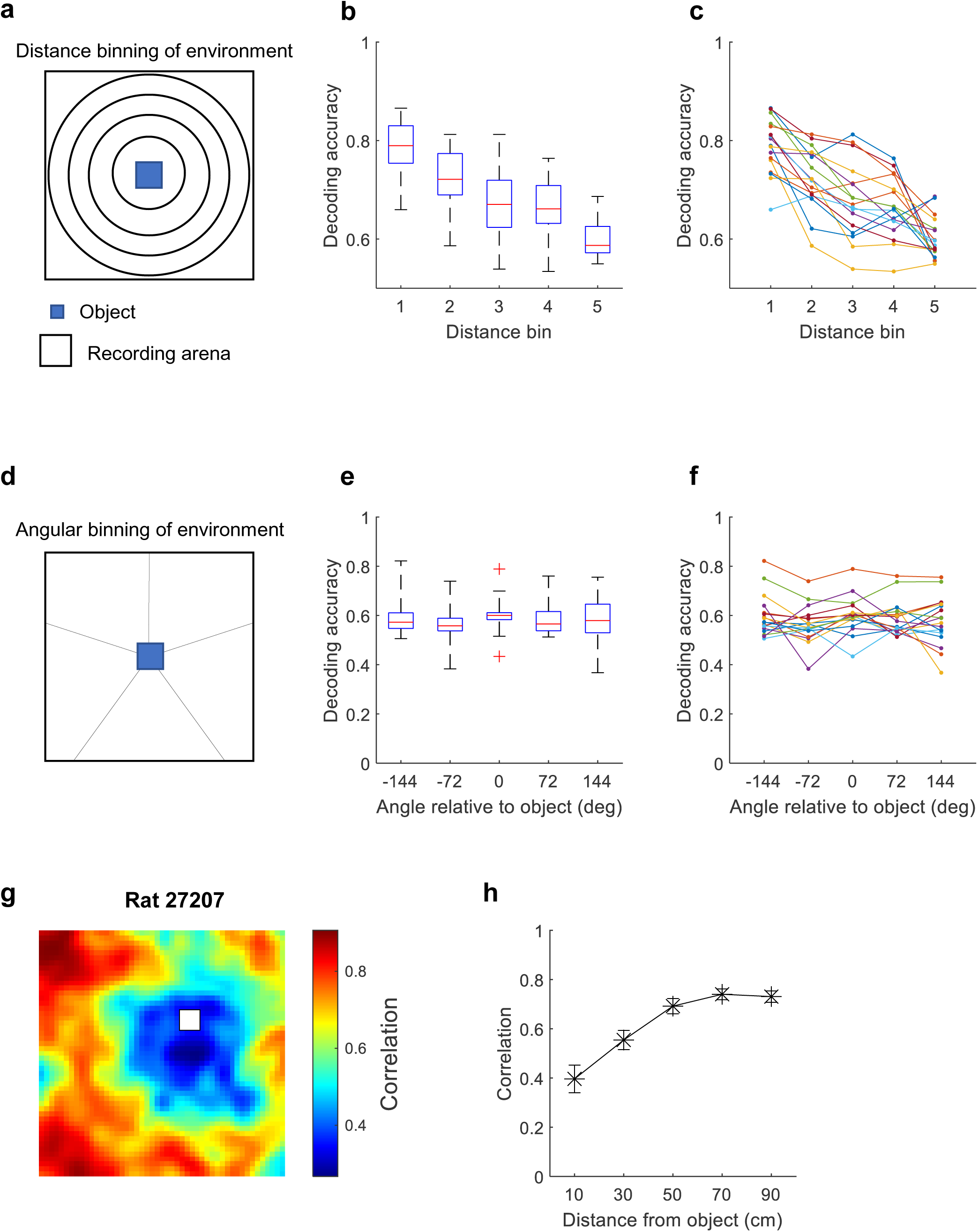
Graded reorganisation of hippocampal population activity. **a,** Schematic of partitioning of the environment for the main analysis. We divided the recording arena (150 cm x 150 cm) into 5 distance bins, forming circles around the object. We then applied the decoder to data from each distance bin to assess how decoding accuracy scales with distance from object. **b**, Box plot of decoding accuracy as a function of distance from object. The central dashed line indicates the median, and the top and bottom lines indicate upper and lower and quartiles, respectively. The length of the whiskers indicates 1.5 times the interquartile range. Outliers exceeding 1.5 times the interquartile range are shown as dots. Distance bin 1 is nearest the object; distance bin 5 furthest away. Note the steady decrease in decoding accuracy as a function of distance from object, suggesting that neural population activity is sensitive to the presence of an object and that this sensitivity is distance-dependent. **c,** Decoding accuracy as a function of distance from object for each individual experiment (n = 17). Data are the same as in the previous panel, but each colour shows a different experimental session. Note that all experimental sessions have decreasing curves. **d,** Schematic of the partitioning of the environment for angular binning. We divided the arena into 5 angular bins, each bin spanning 72°. Then, we applied a decoder to each bin to assess how decoding accuracy scales with angle relative to object. **e,** Box plot of decoding accuracy as a function of angle relative to object. **f,** Decoding accuracy as a function of angle relative to object for each individual experiment. Data are the same as in the previous panel, but each colour shows a different experimental session. Note that all experimental sessions have flat curves. **g**, Correlation map showing similarity of neural activity on trials with and without the object (rat 27207). We obtained the correlation map by taking the two stacks of rate maps from the No Object trial and Object trial and computing their pairwise Pearson correlation. This gives a correlation value (from −1 to 1) for each spatial bin, indicating the amount of correlation across different regions of space. Note that correlations are closer to zero (near 0.3) in the vicinity of the object and more positive (beyond 0.8) in the periphery. **h,** Correlation between stacks of rate maps (No Object trial vs. Object trial) as a function of distance from the object. For each experimental session, we calculated a correlation map as in the previous panel. Then, we binned all values in the correlation map by distance to object and calculated the mean correlation inside each distance bin. Plot shows mean ± SEM for all experimental sessions (n = 17) from all animals.

Having shown a reorganisation of population activity as a function of distance from the object, we then wondered if the reorganisation was equally large in all directions relative to the object. To test this idea, we divided the environment into 5 angular bins, with each bin covering a different set of allocentric angles relative to the object (Fig. 4d). The decoding accuracy was the same in different angular bins (Fig. 4e, f) (*H* = 3.75, *p* = 0.4416, Kruskal-Wallis test, n = 17 experimental sessions). This shows that the distance-dependent reorganisation of population activity is general and not specific for certain compass directions.

To ensure that the result reflects a genuine phenomenon in the population activity and is not an artefact of the decoder, we performed several controls. First, we observed no decrease in decoding accuracy across the same distance bins when attempting to decode the first half of the No Object trial from the second half of the No Object trial (Supplementary Fig. 6c, d). This means that, without an object present to induce the reorganisation, we do not observe the graded decrease. Second, we randomly shuffled the trial labels of the spike count vectors (labels being “No Object” and “Object”), destroying the association between spike count vectors and trials. That is, due to shuffling, a spike count vector that occurred in “No Object” might now be labelled “Object”. On the shuffled data, the decoder performed with a mean accuracy of 50% in all distance bins (Supplementary Fig. 6e, f). Third, in the standard version of our decoder, every recording session was split into training and testing data by using the first half of the session for training and the second half for testing (and then vice versa). To confirm that this did not impact the result, we partitioned the data into training and testing in a random manner. The results from this decoder showed the same decline in decoding accuracy as a function of distance from the object (Supplementary Fig. 6g, h). Finally, a decoder that used the differences in the amount of time the animal spent in each spatial bin (occupancy) between the No Object and Object trials to infer the presence or absence of the object did not show a distance-dependent decrease in decoding accuracy (Supplementary Fig. 6i, j), suggesting that differences in occupancy across different distance bins do not explain the result.

Since the decoder behind the main result (Fig. 4b, c) used rate maps of hippocampal cells as training data (see Methods for details), we thought that perhaps the population of rate maps provides a more striking view of the hippocampal object representation than rate maps of single cells. Therefore, we took the entire stack of rate maps from the No Object trial and Object trial and computed the pairwise Pearson correlation between the two stacks. This gives a correlation value (from −1 to 1) for each spatial bin (size: 3 cm×3 cm), showing how correlated the two stacks of rate maps are at each location. The correlation maps had low correlations in the central regions near the object, but high correlations in peripheral regions further away from the object (Fig. 4g), indicating that the neural population activity reorganizes as a function of the animal’s proximity to the object. Next, we binned the environment into 5 distance bins (using fixed bins of 0-20 cm, 20-40 cm, 40-60 cm, 60-80 cm and 80-100 cm) and collected the correlation values in each bin (number of correlation values for each bin: 489 ± 27; mean ± SEM; n = 17 experimental sessions). This showed a stepwise increase in correlation values as a function of distance from object (Fig. 4h) (*H* = 30.1, *p* = 4.67×10^−6^, Kruskal-Wallis test; significantly different correlation values for distance bins 1 vs.2; 2 vs. 3; and 3 vs. 4, *W* ≤ 19, *p* ≤ 0.0113, Wilcoxon signed-rank tests). Altogether, these findings suggest that the object acts as a pivot around which the hippocampal population activity reorganizes: In the vicinity of the object, the changes in the population activity are large, producing high decoding accuracy (and low correlations), whereas further away, the changes are small, producing low decoding accuracy (and high correlations).

### Objects are encoded by distinct patterns of joint activity in CA1 cells

Having shown that the hippocampal population activity contains information that can be used to decode the presence of an object in the animal’s environment, we next asked: can we identify specific features of the population activity that become different when the object is introduced? Given the results of the decoder (Fig. 4b, c), we are looking for a specific feature of the population activity that (1) is different between the No Object and Object trials and (2) becomes more and more different with proximity to the object. To parametrize the population activity, we will introduce the concepts of “words” and “counts” as used previously to study neural coding at the population level^27–29^. We define a word as a unique pattern of spiking and silence among all neurons in the population in a single time bin (time bins of 10 ms; see Methods for rationale behind this choice). For example, in a population of four neurons, [0, 0, 0, 1] is a unique word and [1, 0, 0, 1] is another unique word because the patterns of spiking and silence are not identical (Fig. 5a). Words provide a detailed description of the population activity that preserves information about neuronal identity (which neurons fired which spikes). In contrast to this description, we define “counts” as the total sum of spikes across the population in a single time bin. For example, [0, 0, 0, 1] contains a total count of 1 and [1, 0, 0, 1] contains a total count of 2. Counts provide a coarse description of the population activity that lacks information about neuronal identity.

**Figure 5.**
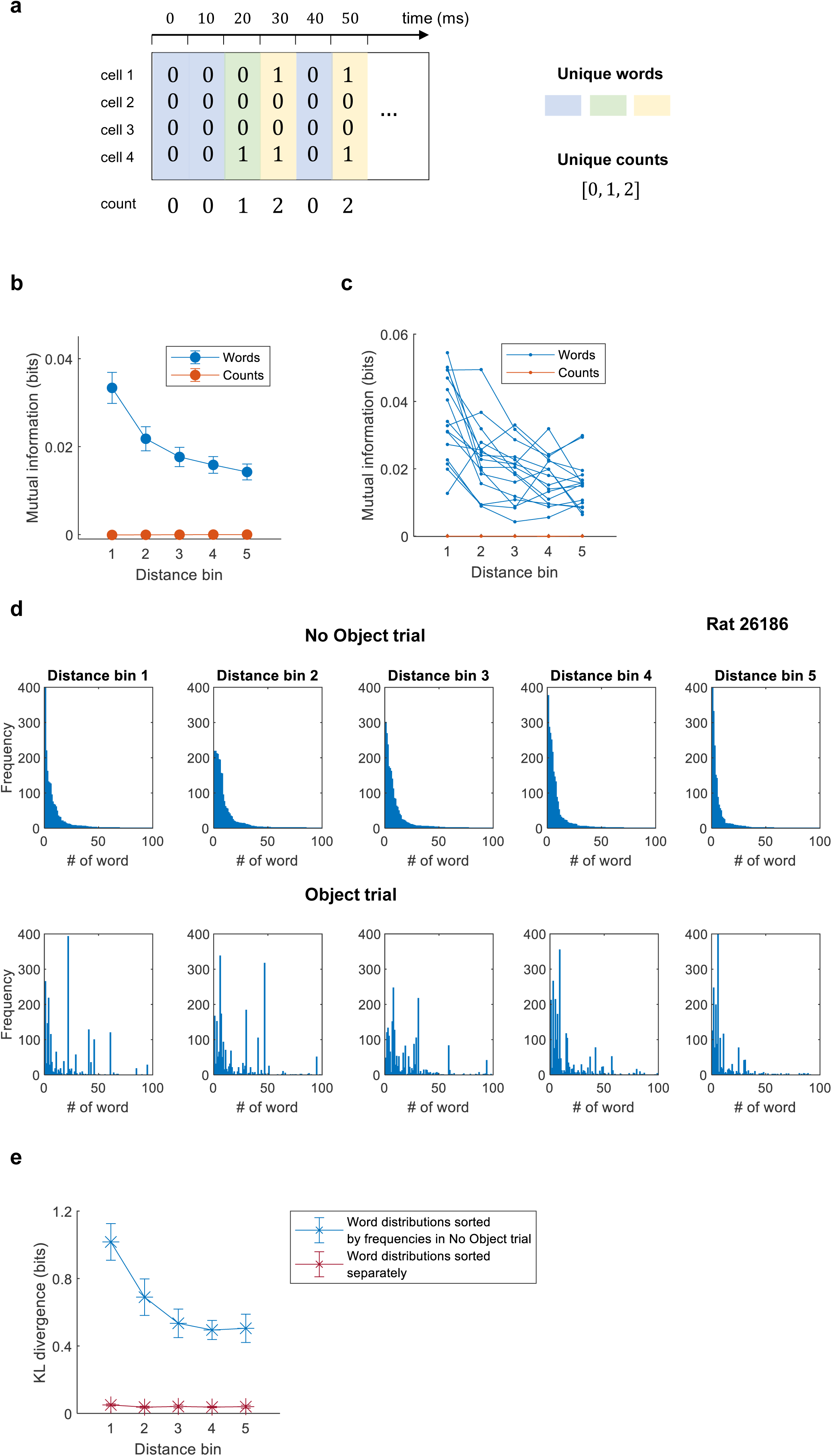
Objects are represented by distinct patterns of joint activity in the CA1 cell population. **a,** Framework for analysing population activity. For population analyses, we represent neural data in an N×T spike matrix, where N is the number of recorded cells and T is the number of timepoints in the trial. Each entry of the matrix contains the number of spikes generated by each cell in each time bin. Using 10 ms time bins we obtain mostly ones and zeroes (in rare cases, 2 or 3). We define a specific pattern of spiking across the neural population as a “word”. In the schematic, each unique word is labelled by a different colour. Words tell us about the detailed spiking pattern across the neural population. We also define “counts” as the total number of spikes in the neural population in any given time bin. The total count, illustrated in the bottom row, is a coarser description of population activity, lacking information about which neuron fired which spikes. **b,** Mutual information between trial type (No Object or Object) and population response (either words or counts) as a function of distance from object. The mutual information quantifies how much information we gain about the trial type by observing words (blue) vs. counts (orange). Distance bins were constructed to contain the same number of samples, as in Fig. 4a, b, c. Data points show mean ± SEM (n = 17 experimental sessions). Distance bin 1 is nearest the object; distance bin 5 furthest away. Words but not counts are informative about trial type, and they are more informative if activity is sampled near the object. **c,** Mutual information from words (blue) or counts (orange) for each individual experiment. Data and analysis are as in the previous panel, but each curve shows a different experimental session. Notice that all orange curves overlap. **d,** Histograms of word frequencies from the No Object trial (top) and the Object trial (bottom). Each histogram shows how many times each unique word appeared during each trial, in each distance bin. The words are indexed from 1-100 (only the 100 most frequent words in the No Object trial are plotted). All distributions are sorted according to how frequently each word occurred in the No Object trial. For this reason, distributions from the No Object trial are decreasing (top) while distributions from the Object trial have more arbitrary shapes (bottom). This also means that indices of words match between top and bottom (word #1 at the top is the same as word #1 at the bottom). **e,** Kullback-Leibler (KL) divergence between word distributions as a function of distance bin. The KL divergence, in this case, quantifies how much each Object distribution (Fig. 5d, bottom) differs from its corresponding No Object distribution (Fig. 5d, top). Word distributions were either sorted by word frequencies in the No Object trial (as described in the previous panel, Fig. 5d) or independently (by word frequencies within the same trial). The former sorting enabled us to assess if frequencies of the same words changed (blue). The latter (independent) sorting enabled us to assess if, for example, one trial type used a larger number of words (of low frequency) and the other trial type used a smaller number of words (of high frequency).

Equipped with these two descriptions of population activity, we wanted to compare their relative merit in explaining the results of the decoder (Fig. 4b and c). Therefore, we calculated the mutual information (MI) between the words and the trial (No Object and Object), telling us how much information we gain about the trial (i.e., whether the object is present or not) by observing the words that the hippocampus generated:

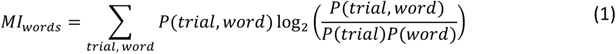

Similarly, we then calculated how much information we gain about the trial by observing the counts:

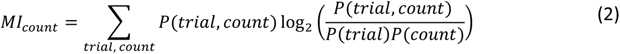

To account for the well-known problem of overestimating MI on finite datasets^30^, we calculated the amount of information expected by chance only due to finite sampling, and subtracted that amount from the MI calculated on real data (see Method for details).

Words were much more informative than counts (Fig. 5b) (MI between words and trials was significantly larger than MI between counts and trials for all distance bins: *U* ≥ 439, *p* ≤ 1.19×10^−6^, Wilcoxon rank-sum tests).Importantly, the amount of information gained from the words decreased as a function of distance from the object (Fig. 5b) (*H* = 24.29, *p* = 6.98×10^−5^, Kruskal-Wallis test), reproducing the pattern of the decoder. The fall-off in mutual information as a function of distance relative to the object was also evident in individual experiments (Fig. 5c). In contrast, the amount of information gained from the counts did not depend on distance from the object (*H* = 4.08, *p* = 0.3947, Kruskal-Wallis test) and was not significantly different from zero (*t* ≤ 1.46, *p* ≥ 0.1636 for all comparisons, t-tests). This means that words, and not counts, underlie the distance-dependent reorganisation of the population activity around the object.

To directly visualize the object-induced change in word representation, we identified the unique words produced by the hippocampus in each experimental trial and calculated their frequencies. We then sorted the words according to how frequently they occurred in the No Object trial (Fig. 5d, top). By using the same sorting of the words in the Object trial (Fig. 5d, bottom) we could directly see which words were up- and down-regulated with an object present. We observed many low-frequency words in the No Object trial occurring with high frequency in the Object trial, standing out as spikes in the distribution (Fig. 5d, bottom). No such spikes were present in shuffled versions of the data (Supplementary Fig. 7a, c), suggesting that the frequencies of specific words changed and that No Object and Object trials recruited different patterns of neural activity.

Looking at the change in word frequencies across distance bins, distributions were very different near the object (Fig. 5d, distance bin 1) but became increasingly similar with further distance from the object (Fig. 5d, distance bins 2-5). To quantify the dissimilarity between the word distributions from the No Object and Object trials, we calculated the Kullback-Leibler (KL) divergence (the larger the KL divergence the more different the distributions are). Word distributions had larger KL divergences near the object and smaller KL divergences further away (Fig. 5e, blue curve) (*H* = 16.22, *p* = 0.0027, Kruskal-Wallis test; KL divergences in 1st distance bin – 5th distance bin: *W* = 102, *p* = 6.10×10^−4^, Wilcoxon signed-rank test), meaning that word distributions became increasingly trial-specific as the distance between animal and object decreased.

While this shows that the hippocampus uses specific words with different frequencies in the No Object and Object trials (sometimes involving different words when the count of a word is changed from *count = 0* to *count > 0*), we wondered if the word distributions would still be different if they were sorted independently (rather than sorted in the same way). This would be the case, for example, if one trial type used a small number of words (occurring with high frequency) and the other trial type used a large number of words (occurring with low frequency). We therefore sorted word distributions from both trials separately (in order of descending frequency in each trial). With independent sorting, the word distributions appeared identical across all distance bins (Supplementary Fig. 7b) and the KL divergence was small and flat (Fig. 5e, red curve) (mean values: 0.0503, 0.0373, 0.0410, 0.0377 and 0.0400; *H* = 3.35, *p* = 0.5004, Kruskal-Wallis test). Direct calculation confirmed that the number of words was similar across distance bins (Supplementary Fig. 7d, e) and between the No Object and Object trials (*z* = 0.221, *p =* 0.8252, Wilcoxon rank sum test). This means that the hippocampus neither uses more nor fewer words to represent the object. Rather, it is the frequencies of specific words that differ between the trials (Fig. 5d, e). Thus, when an object is present in the environment, we observe that (1) the hippocampus changes its use of unique activity patterns (upregulating some patterns and downregulating others) compared to when no object is present and (2) the recruitment of activity patterns becomes increasingly different as a function of the animal’s proximity to the object.

### Distance-specific reorganization of the hippocampal map is widely distributed across the CA1 population

We next wondered how widely distributed the distance-based reorganisation of the hippocampal map is across CA1 cells. Note that we use the term “distributed” in a functional sense rather than an anatomical sense (i.e., we ask how widely spread the reorganisation is across the recorded CA1 cells). One possibility is that the reorganisation is driven by a dedicated subpopulation of cells with “strong tuning” to distance from the object. Alternatively, the reorganisation could be widely distributed across the recorded CA1 cells, so that it is an emergent property of the network and largely independent of how the responses look at the single-cell level. To distinguish between these possibilities, we asked: (1) How many cells do we need to see the reorganisation? (2) Can we see the reorganisation in the joint activity of cells that are weakly tuned to distance? (3) Are specific single-cell properties (for example, high object-vector score or high spatial information) predictive of a larger contribution to the reorganisation?

To understand how many cells are necessary to see the reorganisation (question 1), we first asked if it is present already at the level of single cells. To estimate how much the cell’s response is modulated by distance from the object, which we refer to as their ‘distance tuning’, we first ran the decoder (Fig. 3a) using only one cell at a time (n = 1189 cells and runs of the decoder). When pooled together, individual cells showed no obvious decrease in decoding accuracy as a function of distance, with medians just slightly above 0.5 (Fig. 6a, compare to Fig. 4b). We then calculated

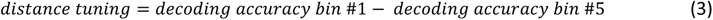

as a proxy for distance tuning of individual cells, where a more positive value implies stronger distance tuning. CA1 cells showed a wide distribution of distance tuning values (Fig. 6b), with no evidence for a strongly tuned subset of cells. There was only a slight bias towards positive values (Fig 6b, white stippled line), consistent with the distance reorganisation not being strongly expressed at the level of single cells.

**Figure 6.**
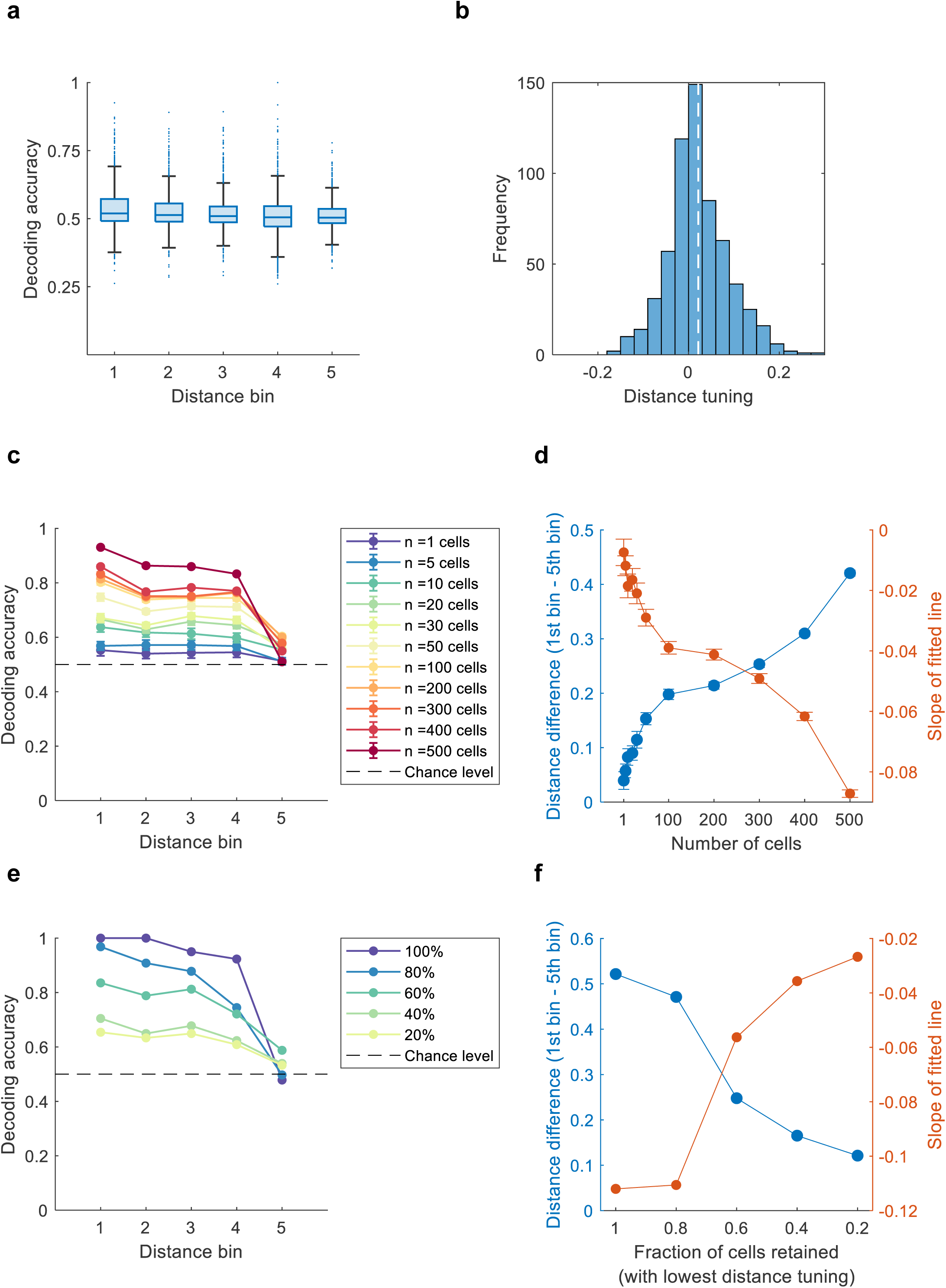
The distance-specific reorganisation is widely distributed. **a,** Decoding accuracy as a function of object distance for individual cells. All cells used for population analyses (15 tetrode sessions and 2 Neuropixels sessions, n = 1189 cells) were fed into the decoder, one cell at a time. Individual cells have mostly flat decoding accuracy. Box plot conventions are as in Figure 4. **b,** Distance tuning of individual cells (n = 1189 cells). After running the decoder with individual cells, for each cell we calculated its distance tuning as the difference in decoding accuracy between bin #1 and bin #5. White stippled line shows mean. Note wide distribution of distance tuning values, as well as the bias towards positive values. **c,** Decoding accuracy as a function of distance from object, depending on how many cells are used in the decoding. Data were generated by taking a random subset of n cells from a total population of N = 620 cells and then running the decoder. The number of cells was varied between n = 1, 5, 20, 30, 50, 100, 200, 300, 400 and 500. For each specific n, we repeated the process 20 times, using different subsets of cells on each iteration. Data points show means; error bars show SEM. When error bars cannot be seen it is because they are too small. Data are from a rat implanted bilaterally in the hippocampus with Neuropixels probes. Stippled line indicates chance level. **d,** Distance reorganisation as a function of number of cells used in the decoding. The left y-axis (blue curve) shows the distance difference (decoding accuracy in bin #1 – decoding accuracy in bin #5) and the right y-axis (orange curve) shows the slope of the least-square line (fitted to decoding accuracy vs. distance curves in the previous panel). These two measures are proxies for the amount of distance reorganisation. Datapoints show means; error bars show SEM. Note increasing distance difference, as well as the decreasing slope, as a function of the number of cells. **e**, Decoding accuracy as a function of distance from object, depending on the fraction of cells with lowest distance tuning retained. We calculated distance tuning of single cells (as in Fig. 6a and b), sorted the cells according to their distance tuning, and ran the decoder using the k% of cells with lowest tuning, with k varying between 100%, 80%, 60%, 40% and 20%. Note that keeping 20% of cells with the least tuning is sufficient to see reorganization of neural population activity according to distance from the object. Data are from the same Neuropixels rat as in panels a, b. **f,** Distance reorganisation as a function of the percentage of cells with the lowest distance tuning retained. Conventions are as in panel d.

We next asked how the reorganisation scales with the number of cells used in the decoder (question 1). For this, we used Neuropixels data from a rat implanted with two probes in the left hippocampus, yielding 620 cells with a wide variety of properties, including place cells, non-place cells, interneurons, head-direction cells and small numbers of object cells and object-vector cells. To simulate different population sizes, we picked a random subset of cells (subsets of size *n* = 1, 5, 10, 30, 50, 100, 200, 300, 400, 500) from the entire population of *N* = 620 cells. We then ran the decoder using this subset, repeating the whole process 20 times. This meant using 20 different subsets of cells for each n, with each subset having a mixture of cell identities and properties. All curves showed reduced decoding accuracy as a function of distance, except the curve constructed using only a single cell (Fig. 6c) (n = 1 curve: *W* = 2.48, *p* = 0.6489; all other curves: *W* ≥ 11.89, *p* ≤ 0.0182, Wilcoxon signed-rank tests). Importantly, the difference between bins #1 and #5 grew systematically with the number of cells (Fig. 6d; left y-axis, blue) (*H* = 174.57, *p* = 3.14×10^−32^, Kruskal-Wallis test) and reached its maximum at n = 500 cells. In parallel to this, the slope of the least-square line (fitted to the curves in Fig. 6c) became increasingly negative (Fig. 6d, right y-axis, orange) (*H* = 157.6, *p* = 1.011×10^−28^, Kruskal-Wallis test) and reached its minimum at n = 500 cells, creating the appearance of two curves that were mirror-images of one another. First, this implies that even a small number of cells picked randomly is sufficient to see the distance modulation, independently of the individual cell properties (Fig. 6c, blue and green curves). Second, the more cells we include, the stronger is the distance-modulated reorganisation of the map (Fig. 6d). This means that increased population sizes allow us to visualise the reorganisation more and more clearly, suggesting that the factors that underlie the reorganisation are widely distributed across the entire population of hippocampal cells.

To verify the distributed nature of the code, we next asked whether we can visualise the distance reorganisation even when using cells that show the least ‘distance tuning’ (question 2). Therefore, after calculating the distance tuning of single cells (Fig. 6a and b) (equation 3), we sorted the cells according to this variable, starting with the lowest value. We then subsampled the same Neuropixels data as before (Fig. 6c, d), keeping either 100%, 80%, 60%, 40% or 20% of cells with the lowest distance tuning. As expected, curves gradually became flatter when using smaller populations of cells (Fig. 6e). However, even the 20% of cells with lowest tuning was sufficient to demonstrate the distance modulation, in the form of positive distance tuning as well as a negative slope (Fig. 6e, f; rightmost point on blue curve: 0.121; on orange curve: −0.0267). This is a direct demonstration of a widely distributed code, where even the “worst” cells (with lowest distance tuning) contribute to a distance-specific reorganisation of the hippocampal map. Such a distributed population code is analogous to how non-place cells also provide information about an animal’s location in space^18^.

Finally, we asked if any properties of single cells were predictive of distance tuning (question 3). For example, do cells with higher object-vector score (Fig. 1) or higher spatial information have stronger distance tuning than other cells? Using the distance tuning of individual cells (Fig. 6b) (equation 3), we correlated this quantity against the object-vector score, spatial information, spatial correlation, or rate overlap (Fig. 7a-d, insets). Because of the large number of cells in our dataset, we present binned versions of these plots for visualisation purposes (Fig. 7a-d, main panels). Specifically, we binned each single-cell property (using 10 bins, ranging from minimum to maximum observed value) and calculated distance tuning in each bin. Distance tuning did not appear to depend strongly on any single-cell property (Fig. 7a-d). The object-vector score was not significantly correlated with the distance tuning (*r* = 0.0054, *p* = 0.84). Spatial information had a significant negative correlation, but the correlation was very weak (*r* = −0.13, *p* = 3.70×10^−7^). Similarly, we looked for any relevance of spatial correlation or rate overlap (Fig. 2) as measures of how strongly cells change their firing locations or firing rates between the No Object and Object trial. Both were negatively correlated with the distance tuning, but in both cases very weakly so (spatial correlation: *r* = 0.08, *p* = 7.61×10^−4^; rate overlap: *r* = −0.18, *p* = 1.57×10^−9^). We obtained similar results when using the slope of the line as a measure of reorganisation (as in orange curves in Fig. 6d, f) rather than distance tuning (object-vector score: *r* = 0.006, *p* = 0.82602; spatial information: *r* = 0.10, *p* = 3.58×10^−5^; spatial correlation: *r* = −0.08, *p* = 7.98×10^−4^; rate overlap: *r =* 0.19, *p* = 1.13×10^−12^). Given the lack of even moderate correlations, it appears that cells can contribute to the distance fall-off independent of whether they are object-vector cells or not (Fig. 7a), place cells or non-place cells (Fig. 7b) or have small or large changes in spatial selectivity (Fig. 7c) or in firing rate (Fig. 7d). The dissociation of our findings from single-cell properties (Fig. 7) is supportive of the notion that distance-dependence is widespread throughout CA1 cells and manifests at the neural population level (Fig. 6).

**Figure 7.**
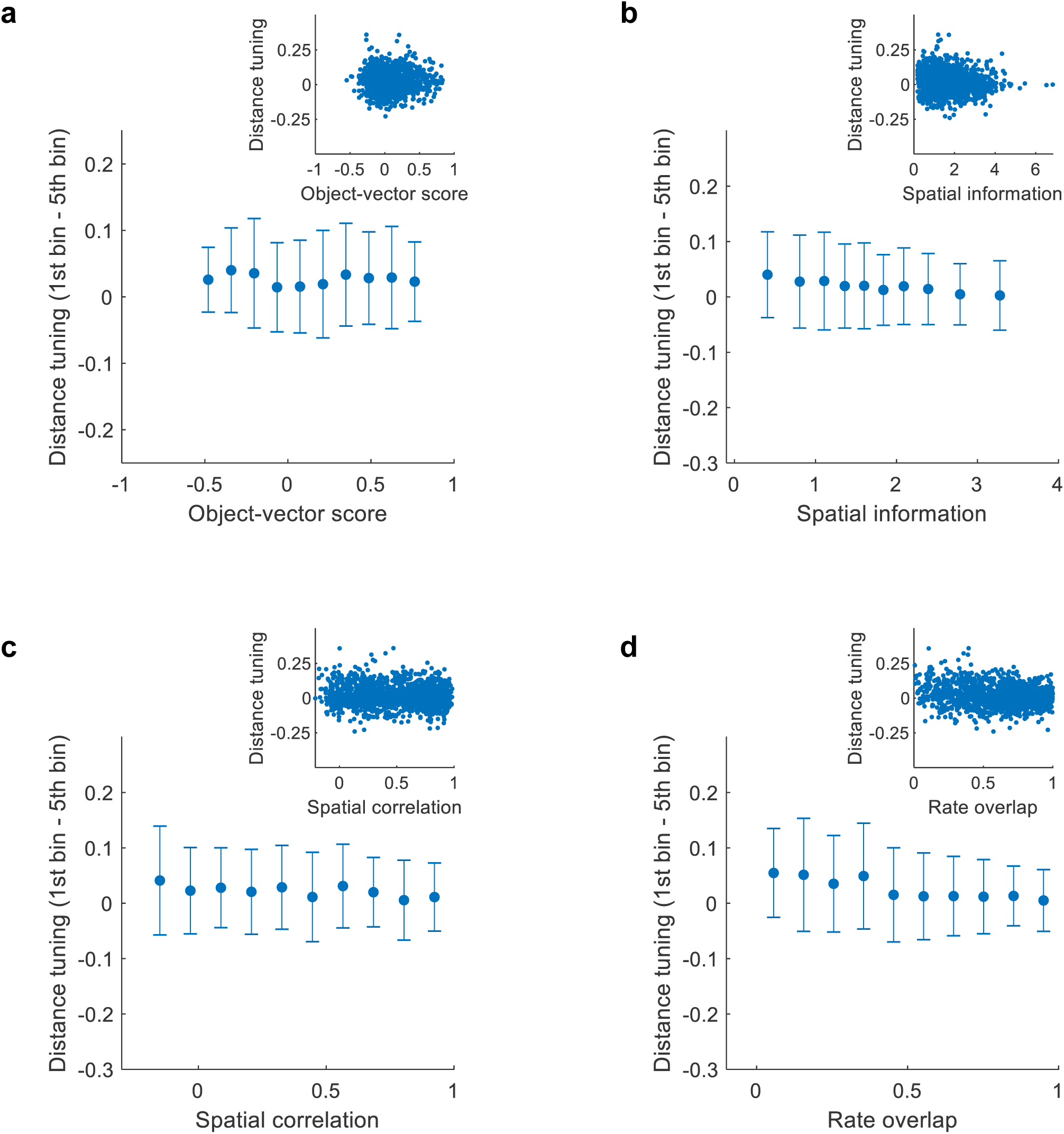
Single-cell properties are not predictive of distance tuning. **a,** Relationship between the object-vector score and distance tuning of individual cells. Distance tuning was estimated by running the decoder with individual cells, one at a time, and then calculating (decoding accuracy in bin #1 – decoding accuracy in bin #5) (as in Fig. 6a, b, d, f). Due to the large number of cells, we bin the data to aid visualization (10 bins, ranging from minimum object-vector score to maximum object-vector score). Inset shows scatterplot of original data before binning (n = 1189 cells). The object-vector score is the Pearson correlation between ‘vector maps’ (Fig. 1e). Note absence of relationship between object-vector score and the distance tuning. **b,** Relationship between spatial information (calculated on Object trial) and distance tuning of individual cells (n = 1189 cells). Conventions are as in panel a. **c,** Relationship between spatial correlation (No Object vs. Object trial) and distance tuning of individual cells (n = 1189 cells). Conventions are as in panel a. **c,** Relationship between rate overlap (No Object vs. Object trial) and distance tuning of individual cells (n = 1189 cells). In all panels, note the absence of relationships between single-cell properties and tuning to distance from the object.

## DISCUSSION

By simultaneously recording up to more than 600 principal neurons in hippocampal area CA1 in freely moving rats, we have described how objects in the local environment influence the firing of both individual cells and neural populations. We only found a small fraction of cells that fired at the object (object cells) or at a fixed distance and direction from it (object-vector cells, also called landmark-vector cells^13^). However, the majority of the CA1 cells changed their spatial firing patterns when the object was introduced, moved, or taken out from the environment. The changes took a variety of forms. At the level of neural populations, these heterogenous responses were coherently organised across cells so that the amount of reorganisation of the hippocampal map was not uniform across space but dependent on distance from the object in a graded manner. The reorganisation of the hippocampal map was widely distributed across the hippocampal network and did not require inclusion of specific subsets of cells. This suggests the existence of population coding based on reorganization of activity around salient landmarks, so that the population activity as a whole communicates an animal’s spatial relationship to those landmarks. Our findings hint that some aspects of hippocampal cognitive maps may not be understood through properties of dedicated functional cell types – such as object cells, object-vector cells, or place cells – but rather through emergent properties of cell populations whose activities appear random and disorganized when viewed individually.

We first set out to estimate how many object cells and object-vector cells are present in CA1 when animals explore 2-dimensional spaces, in a dataset where the number of cells is an order of magnitude larger than in previous work^5,10,13^. We find both object cells and object-vector cells only in small proportions (1.9% and 7.1%, respectively), consistent with earlier work^5,10,13^. The present findings vary somewhat from a recent study that recorded CA1 cells in mice running on a treadmill and identified close to 15% of the cells as object cells or landmark-vector cells^14^. However, in that study, the classification of object cells and landmark-vector cells was made without moving the object, which may have increased the estimated numbers of cells with object-related firing. Here, we report only small numbers of cells firing at objects or at fixed positions relative to objects but we cannot exclude that the numbers would be higher in other contexts or behavioural tasks.

The CA1 cells that fire at fixed distances and directions from objects might superficially appear like object-vector cells in MEC of mice^11^. However, the CA1 cells had variable tuning across time and environments, unlike object-vector cells in MEC which are stable over weeks and across environments^11,31^. A different type of “instability” across time was reported in hippocampal landmark-vector cells, which only began to fire at fixed distances and directions from objects once the rat had gained experience with the objects^13^. In the present study we did not observe such an experience-dependent effect. There was no correlation between the rat’s experience and the prevalence of object-vector cells. On the other hand, given the instability of the object-vector responses, we cannot exclude that slow emergence of vectorial firing would be seen in some cells. Taken together, however, the low number of object-vector cells, and the even lower number of object cells, as well as the apparent instability of these cells across time, suggest that they are not the predominant mechanism for object representation in CA1.

Although object cells and object-vector cells were not abundant among individual cells in CA1, the effect that the object had on neural responses in the area was striking. Most cells changed their spatial firing patterns when we introduced the object to the environment (67.7%) or moved it to a new location (69.9%). When we removed the object, a large percentage of cells did not revert to the pre-object firing pattern and many cells showed trace responses (persistence of object-induced firing fields after removal of the object) (21.1%). This agrees with previous studies showing immediate effects of objects on place cells^7,8^ and the fact that CA1 cells can signal the past position of an object^20,21,25^. Although CA1 cells are known to dynamically change activity across time^23,24^, this could not explain the striking changes in spatial firing patterns in the object task, as the changes were much larger than drift over time in control trials without the object.

How exactly did objects influence the spatial firing patterns? Our findings are consistent with studies that report remarkable heterogeneity of object responses in the hippocampus^7,15^. We observed a similarly wide range of object responses, including appearance and disappearance of firing fields, movement of firing fields, object cells, object-vector cells and trace responses. This heterogeneity points to a transformation of the object-centred spatial codes within the entorhinal-hippocampal circuit, where the representation of space is changed from a low-dimensional code in MEC with coherence between cells’ responses across environments^32–34^ to a higher-dimensional code in the hippocampus that orthogonalizes distinct representations of space^20,35,36^. The high dimensionality of hippocampal responses is closely linked to the phenomenon of global remapping, where distinct place-field codes are created for distinct environments^4,37,38^.

In parallel with the large variability of responses at the level of single hippocampal cells, we observed a striking organisation of the cells’ collective activity. At the neural population level, the object acted as a pivot for reorganizing hippocampal activity patterns: in the object’s neighborhood, the reorganization was large, leading to high decoding accuracy, whereas further away from the object, the reorganization became increasingly smaller, leading to low decoding accuracy. Thus, the highly variable activity of the CA1 cells was brought together into a unified population code, where activity was largely determined by the state of a single variable: the animal’s distance from the object. In that sense, the population response had a simple and predictable structure, in comparison with the disorganized activities of individual cells. The graded reorganization of the population response between No Object and Object trials could not be explained by differences in the animal’s occupancy, as we were unable to decode trial type based on occupancy.

The reorganization of population activity is unlike any form of hippocampal place-cell remapping described in the literature^4,37–40^. During global remapping among place cells, the whole population of cells reorganise their spatial firing patterns, providing an entirely new and orthogonal representation of space. Remapping is traditionally thought of as discrete^38,41–43^, and during periods of ambiguity, there may even be discrete flickering between maps^42,44,45^. That type of discreteness differs from the population coding described here, where the amount of reorganization of the hippocampal map is a continuous function of the animal’s distance from the object. The smoothness of the reorganization shows that that the discrete nature of hippocampal maps can break down under some circumstances (e.g., when comparing an environment in the presence and absence of an objects). At the core of this smooth reorganization is the presence of unique activation patterns across the neural population, previously described as words^27–29^, which the hippocampus generates in an increasingly trial-specific manner the closer the animal is to the object.

A critical element of our findings is how widely distributed the object-centered population coding is across the recorded cells. The more cells we put into the decoder, the larger the improvement in decoding accuracy near the object (relative to far from the object). In addition, we could use the 20% with the weakest distance tuning to visualise the distance-based reorganisation. While distance tuning is weak in most individual cells, at the population level the responses of individual cells become transformed into a clear pattern, with a smooth change in the hippocampal map according to the animal’s distance from the object. The transformation has parallels to classic work in the vertebrate retina, where weak pairwise correlations are transformed into strong correlations in population activity^27^, and supports recent work in the hippocampus showing that place cells and non-place cells encode information in a collective and distributed fashion^17,18^. As previously hypothesized^46^, these findings hint that neural representations may often be best described as emergent properties of large numbers of cells.

## AUTHOR CONTRIBUTIONS

A.N., E.I.M., and M.-B.M. designed and planned experiments; A.N. performed the experiments (surgeries, recordings, spike sorting); A.N. performed single cell analyses; S.A. performed neural population analyses; A.N. and S.A. wrote the paper with inputs and edits from all authors; S.G.C., E.I.M. and M.-B.M supervised the project; E.I.M. and M.-B.M. obtained funding.

## ACKNOWLEDGEMENTS

We thank Ø. Høydal and E. Ranheim Skytøen for discussions and R. Gardner for developing the Neuropixels data acquisition and analysis pipeline. We thank A.M. Amundsgård, K. Haugen, K. Jenssen, E. Kråkvik, I. Ulsaker-Janke, and H. Waade for technical assistance. The study was supported by funding from the Research Council of Norway (FRIPRO grant number 300394 to M.-B.M., Centre of Excellence grant number 223262 to M.-B.M. and E.I.M., National Infrastructure grant number 295721 to E.I.M. and M.-B.M); a grant from the K.G. Jebsen Foundation (grant number SKGJ-MED-022); a Synergy Grant to Y.B. and E.I.M. from the European Research Council (‘KILONEURONS’, Grant Agreement No. 951319); the Kavli Foundation (M.-B.M. and E.I.M.); and a direct contribution to M.-B.M. and E.I.M. from the Ministry of Education and Research of Norway.

## FIGURE TITLES AND LEGENDS

**Supplementary Figure 1.**
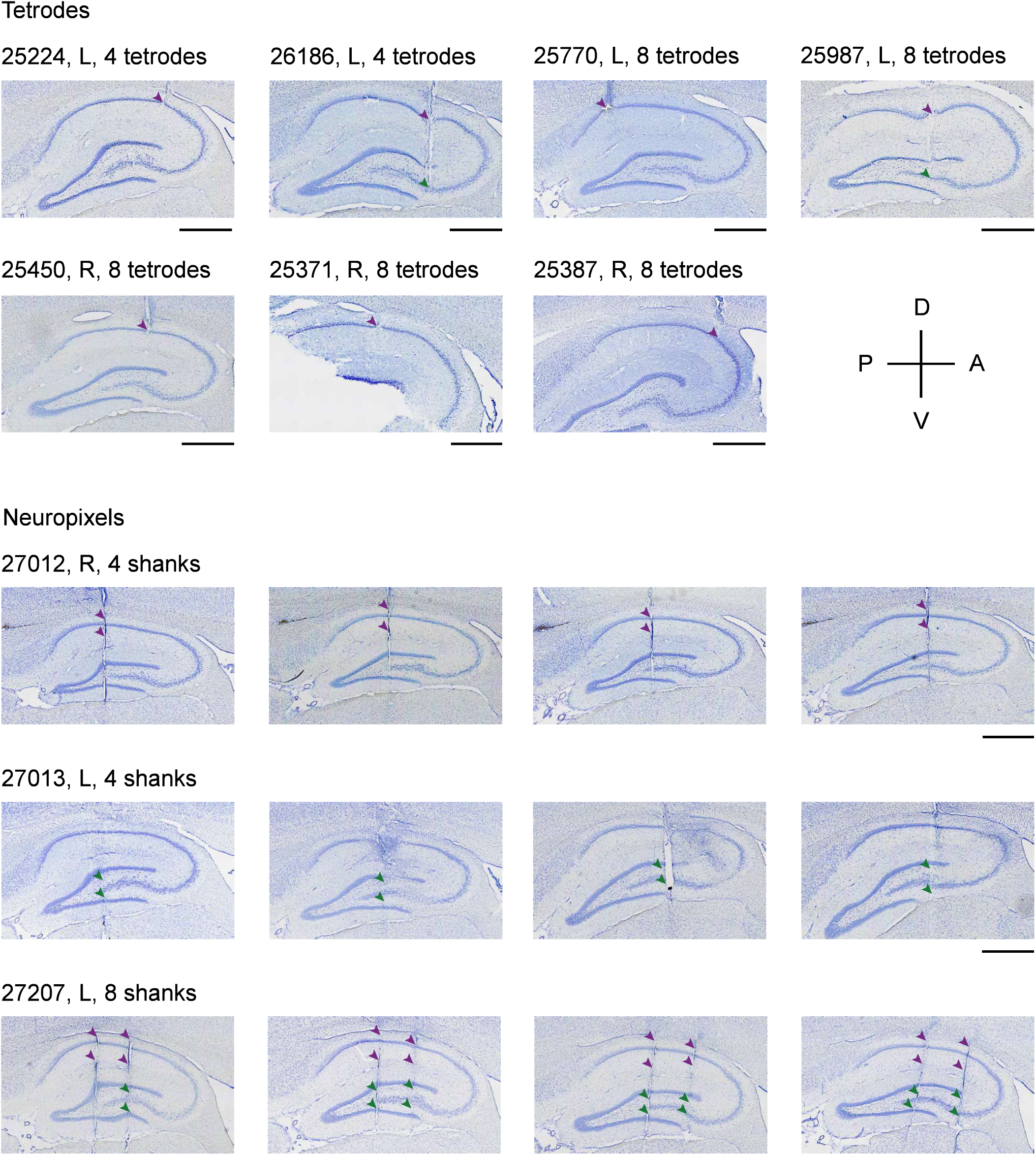
Anatomical location of recording sites. Micrographs of Cresyl violet-stationed sagittal brain sections from 7 rats implanted with microdrives with a single bundle of 4 or 8 tetrodes, and 3 rats implanted with one or two four-shank Neuropixels 2.0 silicone probes. Rat ID, hemisphere (L = left; R = right), and number of tetrodes or shanks is indicated on top of each micrograph. Sections from rats with tetrode implants are organised from most medial (top left) to most lateral (bottom right) implant. Sections from the Neuropixels rats are organised for each rat from the most medially placed shank(s) (left) to the most lateral shank(s) (right). The arrows indicate final recording locations (tetrodes) or top and bottom locations of active recording sites (Neuropixels). Purple arrows indicate CA1 and green arrows indicate CA3/hilus. One brain (rat 25371) was damaged during sectioning, and parts of the hippocampal formation are therefore missing from the micrograph. Scale bars = 1000 µm. D = dorsal; V = ventral; A = anterior; P = posterior.

**Supplementary Figure 2.**
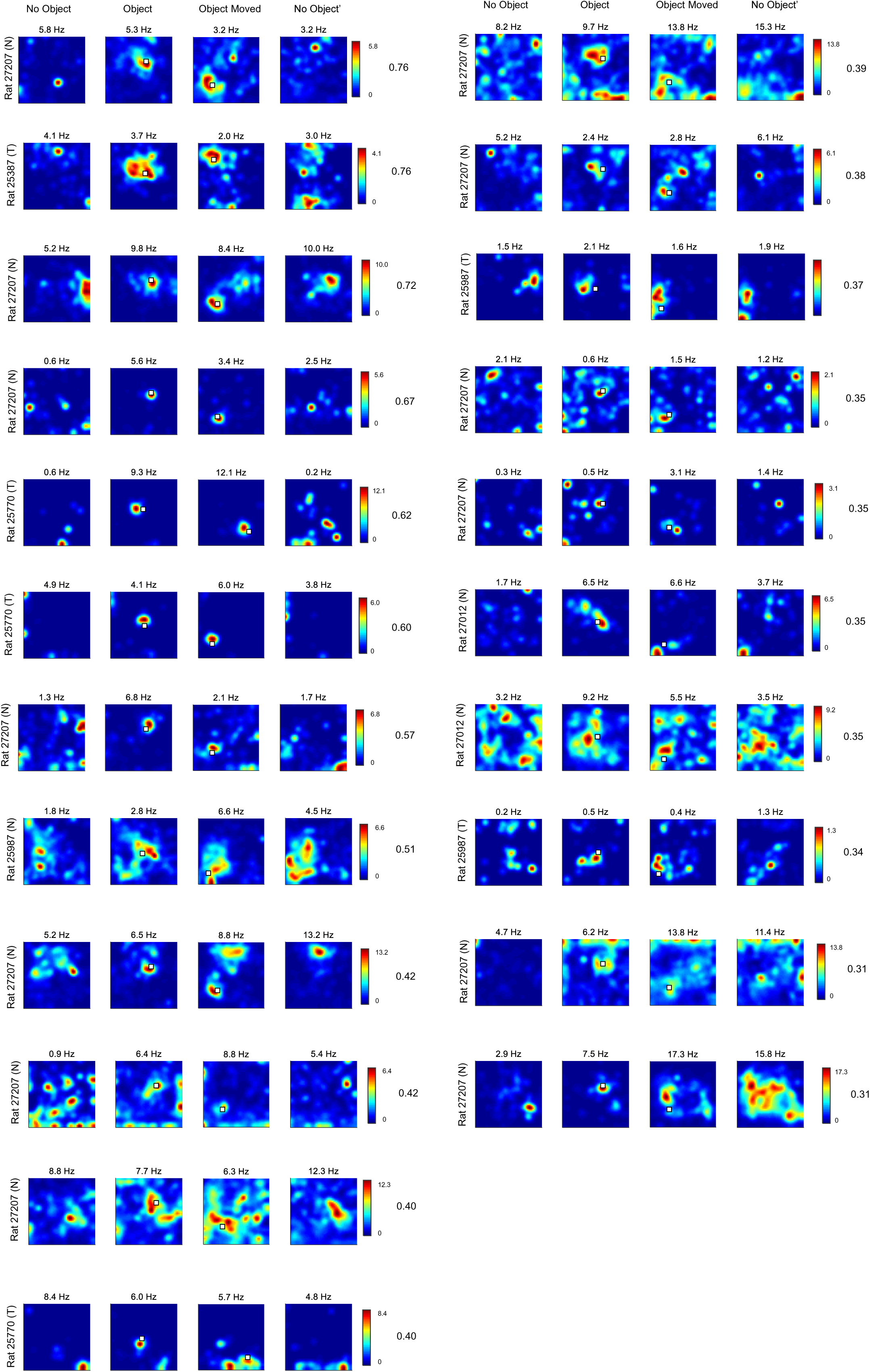
Further examples of object cells. Rate maps for all object cells recorded in CA1 (*n* = 23 cells) from 5 rats. Conventions as in Fig 1b. Object location is indicated by the white square. Cells are ordered according to their object score, starting with the highest. The object score is indicated to the right of each cell. Peak firing rate is indicated at the top of each rate map. Scale bar, colour-coded firing rate. T = rat implanted with tetrodes; N = rat implanted with Neuropixels.

**Supplementary Figure 3.**
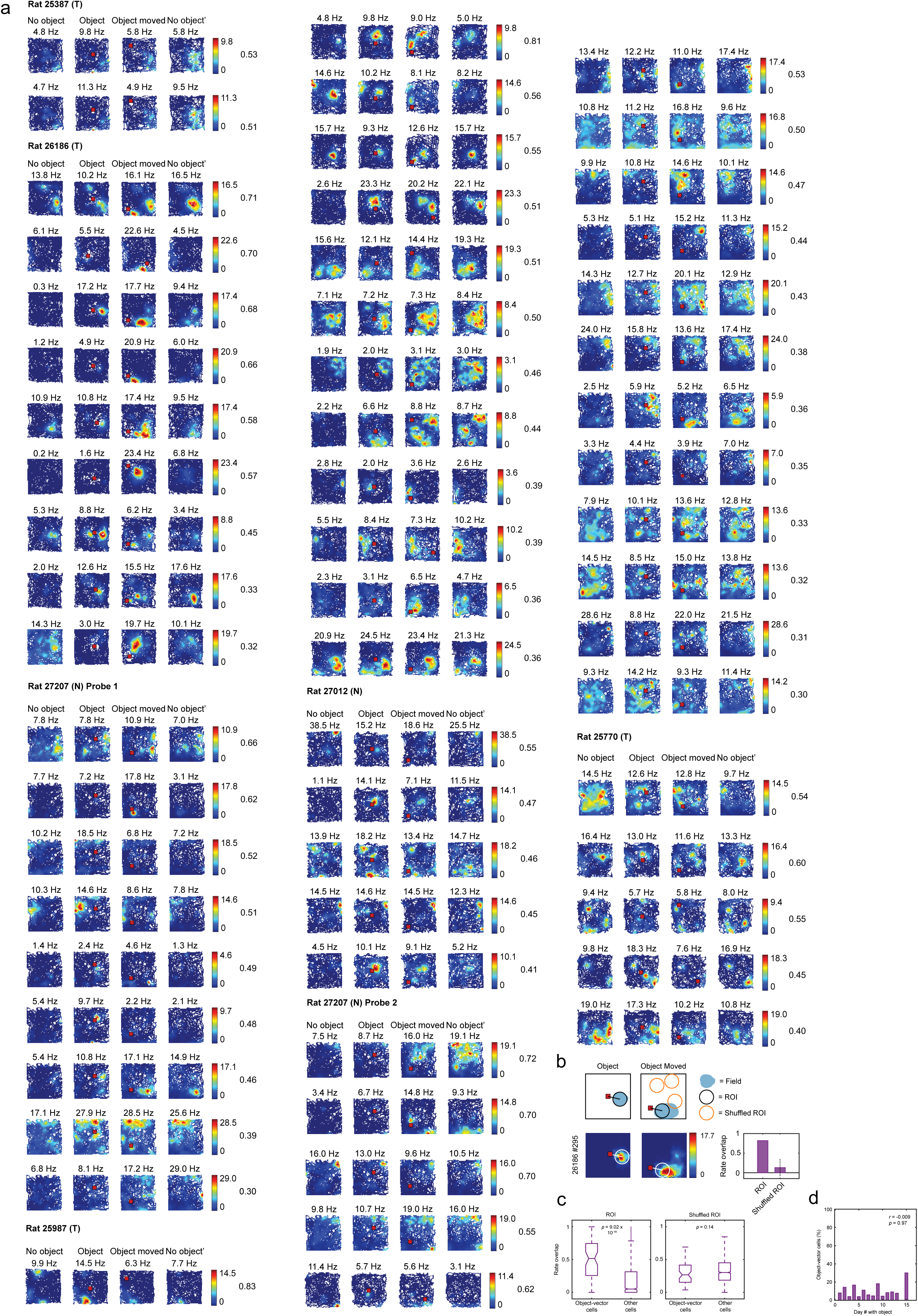
Further examples of object-vector cells and additional information about their properties. **a,** Rate maps for all object-vector cells (*n* = 60 cells) recorded in CA1 from 6 rats. Rate map conventions are as in Supplementary Fig. 2. Rats are organised from most proximal to most distal implant in CA1. For rat 27207, which was implanted with two Neuropixels probes at different proximodistal locations, the cells are separated by probe. **b,** Region-of-interest (ROI) analysis confirms that hippocampal object-vector cells have increased firing rate at fixed distances and directions to objects. Top row, schematic describing the procedure for comparing object-vector fields across object trials. For each cell, in the the firing rate map of the Object trial an ROI (indicated as the black circle) was defined as a circle with centre at the centre-of-mass of the object-vector field (shown in light blue). The vector between the centre of the ROI and the centre of the object (indicated by the red square) was identified. The ROI and the vector were subsequently transferred to the rate map for the Object Moved trial, using the centre of the object in the Object Moved trial as a reference. We then compared the firing rates within the ROIs by calculating the ROI rate overlap, i.e. the firing rate in the less-active ROI (lower firing rate) divided by the firing rate in the more-active ROI (higher firing rate). Then, a shuffling procedure where the ROI was placed at random locations in the Object Moved rate map (‘Shuffled ROI’, indicated in orange) was applied and repeated 1000 times. For each shuffle iteration, we calculated the rate overlap between the ROI in the Object trial, and the Shuffled ROI in Object Moved trial. Bottom left, firing rate maps for an example CA1 cell with ROI indicated as the white circle. Bottom right, ROI rate overlap (0.82) and Shuffled ROI rate overlap (mean ± SD: 0.13 ± 0.21) for the cell to the left. **c.** Left, box plots showing rate overlap between ROIs in the Object and Object Moved trials for object-vector cells (*n* = 60 cells) and for other cells (*n* = 784 cells). For object-vector cells, we expect the rate overlap to be high, whereas it should be low for cells that do not meet the criteria for object-vector cells. Indeed, the rate overlap was significantly different between object-vector cells (*n* = 60 cells; mean ± SEM: 0.50 ± 0.04) and other cells (*n* = 784 cells; mean ± SEM: 0.20 ± 0.01; *z* = 8.04, *p* = 9.02 x 10^−16^, Wilcoxon rank sum test). Right, box plots showing rate overlap between ROI in Object and random placements of the ROI in Object Moved (Shuffled ROI). For each cell, the mean of the 1000 shuffle iterations is used to make the plot. This value was comparable for object-vector cells and other cells (object-vector cells mean ± SEM: 0.29 ± 0.02, other cells mean ± SEM: 0.33 ± 0.01; *z* = 1.44, *p* = 0.15, Wilcoxon rank sum test). **d.** Object-vector score does not depend on experience with the object. Percentages of recorded object-vector cells as a function of day of exposure to the object in CA1 (*n* = 844 cells in 9 rats). We found no significant correlation between the day of exposure to the object and the percentage of recorded object-vector cells (Pearson correlation, *r* = −0.009, d. f. = 14, *p* = 0.97).

**Supplementary Figure 4.**
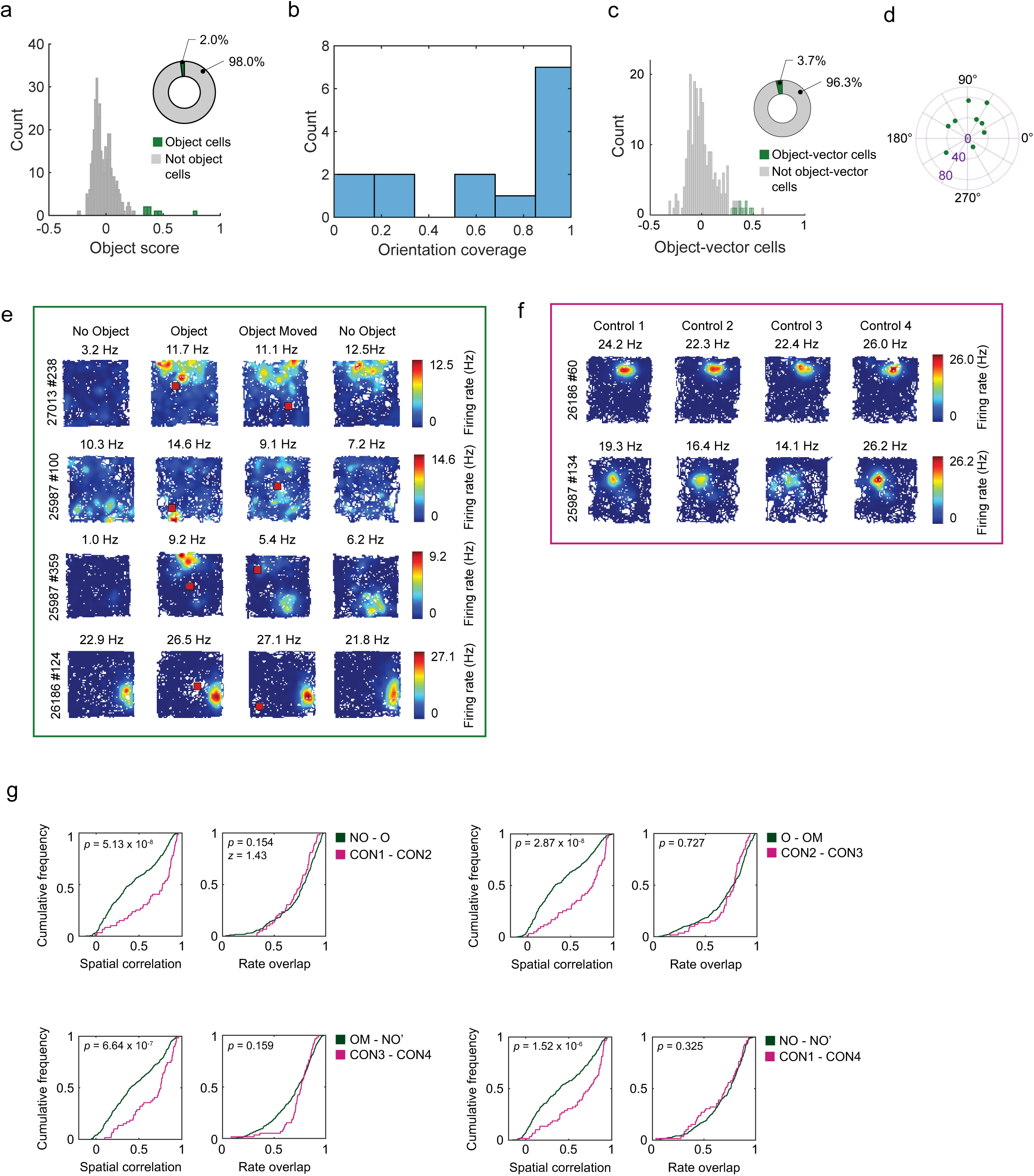
CA3 data corresponding to Figure 1 and 2. **a,** Object cells are very sparse in CA3 and hilus. Left, distribution of object scores for cells recorded in CA3 and hilus. Green bars, cells classified as object cells, grey bars, cells not classified as object cells. Right, pie chart showing percentage of object cells (7 out of 355 cells, 2.0%). **b,** Orientation coverage of object cells in CA3. Object cells mostly fired around the entire object. **c,** Object-vector cells are very sparse in CA3 and hilus. Left, distribution of object-vector scores for cells recorded in CA3 and hilus. Green bars, cells classified as object-vector cells, grey bars, cells not classified as object-vector cells. Right, pie chart showing percentage of object-vector cells (9 out of 243 cells, 3.7%). **d,** Polar scatter plot showing orientation (polar axis, in degrees) and distance (radial axis, in cm, indicated in purple) for object-vector fields in CA3, defined by the centre of mass of the object-vector fields in the vector map. The object-vector fields were located between 19.65-78.76 cm away from the object (mean ± SD: 47.01 ± 18.48 cm) and covered most of the azimuthal range. **e,** Colour-coded firing rate maps for 4 example cells. The top row shows a cell that changed its spatial firing pattern when introducing the object. The second row shows a cell that had a diffuse spatial firing pattern. The third row shows a cell that changed its spatial firing pattern both when introducing and moving the object. The bottom row shows a place cell that was unaffected by the presence of the object. Object locations in the Object and Object Moved trials are shown as red squares. Rat ID and cell ID are indicated to the left of the firing rate maps, and peak firing rate in each trial is indicated at the top of each rate map. Bars, colour-coded firing rate. **f,** We also recorded 59 unique principal cells in 2 rats during control sessions. Colour-coded firing rate maps for 2 example cells. The top row shows a place cell whose peak firing rate remains approximately the same throughout the session, and the bottom row shows a cell where the peak firing rate changes. Symbols as in panel e. **g,** Cumulative normalized frequency distributions of spatial correlation (left plots in each pair) and rate overlap (right plot in each pair). Each plot compares pairs of experimental trials from Object sessions (dark green lines) to the corresponding control trials (pink lines). The spatial correlation is defined as the Pearson correlation between the firing rate maps in two trials, and the rate overlap is calculated by dividing the mean firing rate in the less active trial by the mean firing rate in the more active trial. We calculated differences between the distributions for each panel (dark green lines and pink lines; Wilcoxon rank sum tests; *p*-values are indicated at the top left corner of each plot). As for CA1, we found that spatial correlation values between pairs of trials was always significantly different between Object sessions and control sessions, whereas rate overlap was not significantly different between the two types of sessions. However, the difference was not as pronounced as for CA1, indicating that cells in CA3/hilus are less sensitive to the presence of an object in the environment. NO = No Object; O = Object; OM = Object Moved; NO’ = No Object’; CON1-4 = Control 1-4.

**Supplementary Figure 5.**
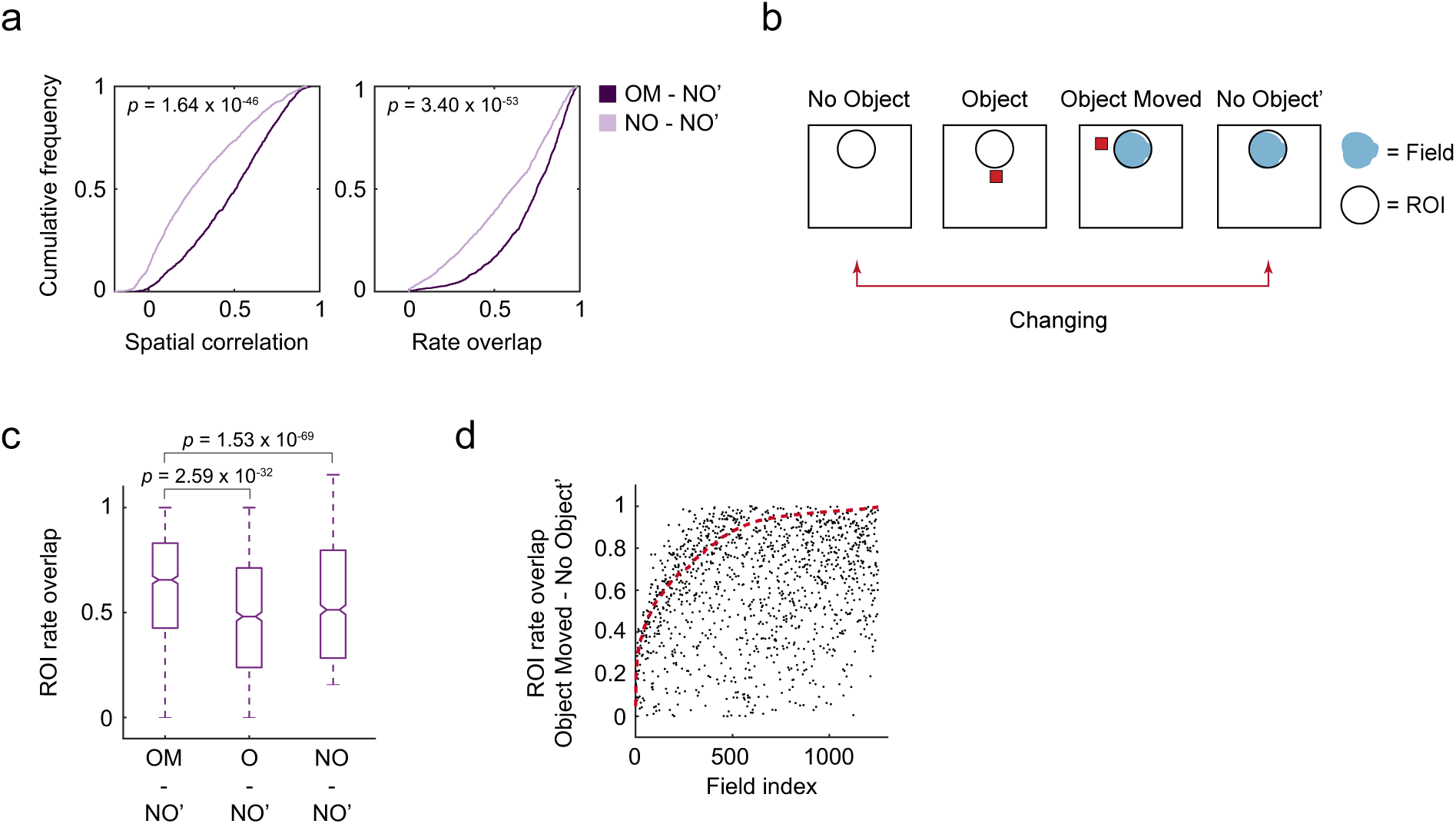
Traces of object-induced firing in CA1 cells. **a.** Normalised cumulative frequency distributions of spatial correlation (left) and rate overlap values (right) between the Object Moved and No Object’ trials (dark purple lines) and between the No Object and No Object’ trials (light purple lines). Both the spatial change (*z* = 14.32, *p* = 1.64 x 10^−46^) and rate change (*z* = 15.35, *p* = 3.40 x 10^−53^, Wilcoxon signed-rank tests) was significantly different between the No Object and No Object’ trials and the Object Moved and No Object’ trials, with higher values of spatial correlation and rate overlap between Object Moved and No Object’ trials. This indicates that many CA1 cells retain their object-induced activity pattern after removal of the object instead of changing back to their pre-object firing pattern. **b.** Schematic showing procedure used to compare the preservation of firing fields across trials. We identified cells where the spatial pattern or firing rate changed between the No Object and No Object’ trials (*n* = 919 cells). In these cells, we identified firing fields (shown in blue) in the No Object’ trial and defined a circular region-of-interest (ROI; indicated by the black circle) around the field. This ROI was then transferred to the corresponding location in the No Object, Object, and Object Moved trials, using the boundaries of the rate map as reference. If the cell had more than one field in the No Object’ trial, we repeated the analysis for each field. The red squares indicate the object. **c.** Box plots showing firing rate overlap between the ROI for *n* = 1,252 fields in *n* = 919 cells in the following pairs of trials: Object Moved and No Object’, Object and No Object’, and No object and No Object’. There was larger rate overlap between the ROIs in the Object Moved and No Object’ trials (mean ± SEM: 0.61 ± 0.01) than between the Object and No Object’ trials (mean ± SEM: 0.48 ± 0.01; *z* = 11.83, *p* = 2.59 x 10^−32^) or between the No Object and No Object’ trials (mean ± SEM: 0.40 ± 0.01; *z* = 17.63, *p* = 1.53 x 10^−69^, Wilcoxon signed-rank tests). This suggests that more cells had a firing field in the same position in the Object Moved and No Object’ trials, indicating carry-over (‘hysteresis’) of firing fields from the most recent object trial. Box plot conventions as in Figure 4b. NO = No Object; O = Object; OM = Object Moved; NO’ = No Object’. **d.** ROI rate overlap values between the No Object’ and Object Moved’ trials for all fields (*n* = 1,252 fields in *n* = 919 cells) plotted as a function of the field index. The red dashed line indicates the 99^th^ percentile of the shuffled distribution for the field (see Fig. 2e, f). Each datapoint represents one field. The fields above the red line passed the 99^th^ percentile of the shuffled distribution and are considered trace fields (*n* = 312/1252 trace fields in *n* = 218/919 cells).

**Supplementary Figure 6.**
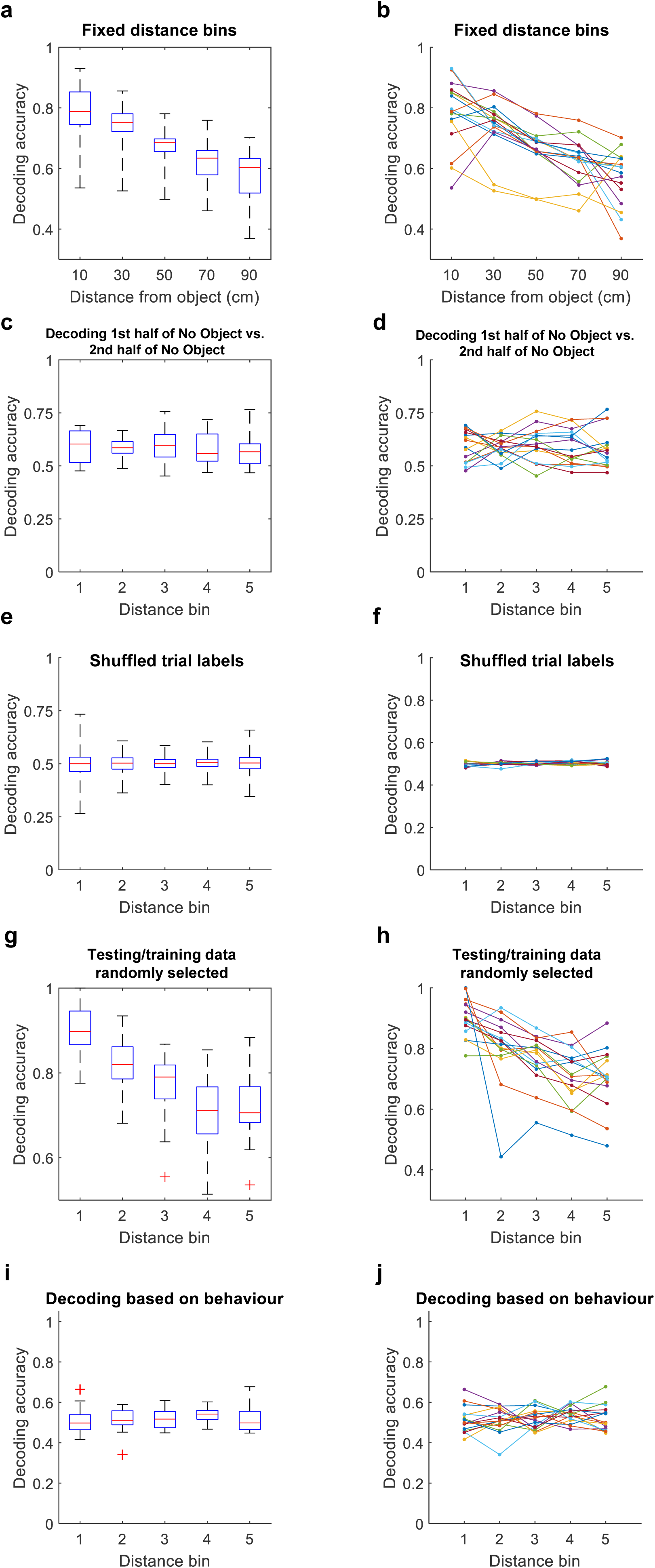
Controls for the results of the decoder. **a,** Decoding accuracy as a function of distance from object when using fixed distance bins. In the main analysis (Fig. 3), we used optimized distance bins to ensure the same number of samples in each bin, avoiding potential biases due to uneven exploration of the animal (for example, if the animal spent little time near the object, we would have few samples in distance bin 1). The fixed distance bins were 0-20 cm, 20-40 cm, 40-60 cm, 60-80 cm and 80-100 cm. Decoding accuracy decreases as a function of distance from object (*H* = 39.69, *p* = 5.02×10^−8^, Kruskal-Wallis test), showing that the result is independent of our choice of binning. **b,** Same data as in the previous panel, but each colour shows a different experimental session. **c,** Decoding accuracy as a function of distance from object when attempting to decode 1st vs. 2nd half of No Object trial (instead of No Object vs. Object). We divided the No Object trial into two halves based on time (for example, if trial was 1800 seconds long, the split was made at 900 seconds). We observed no decrease in accuracy as a function of distance (*H* = 1.46, *p* = 0.834, Kruskal-Wallis test), which is expected given that no object was present in the trial. **d,** Same data as in the previous panel, but each colour shows a different experimental session. **e,** Shuffling trial labels. To ensure that the results were not an artefact of the decoder, we shuffled the trial labels of testing data. This means that a spike count vector was randomly assigned the label “No Object” or “Object” (instead of assigning the label based on which trial it occurred in). The mean decoding accuracy is 50% in all distance bins (mean ± SD in distance bins 1-5: 0.498 ± 0.01; 0.501 ± 0.008; 0.501 ± 0.006; 0.504 ± 0.008; 0.502 ± 0.010; none of the groups significantly different from 0.5: *t* ≤ 2.1254, *p* ≥ 0.0518 for all comparisons, t-tests), confirming that the decoder does not generate distance tuning unless such structure is present in the data. We performed 50 shuffling iterations for each experimental session. **f,** Same data as in the previous panel, but each colour shows a different experimental session. Data are means over 50 shuffling iterations. **g,** Decoding accuracy as a function of distance from object using a random partition of the original data into training and testing data. In the main analysis (Fig. 3), we used the first half of the session as training data and the second half as testing data (and vice versa in a separate iteration). Here, we instead randomly selected half the timepoints to constitute training data. Hence, training data were distributed across the entire session, rather than across one half. The same was true for training data. We observed the same decrease in decoding accuracy as a function of distance from the object (*H* = 39.71, *p* = 4.98×10^−8^, Kruskal-Wallis test), confirming that our results did not depend on the choice of partitioning. **h,** Same data as in the previous panel, but each colour shows a different experimental session. **i,** Decoding accuracy as a function of distance from object for a decoder that infers the trial type (No Object vs. Object) based on the animal’s behaviour (using occupancy maps rather than rate maps). Decoding based on behaviour did not yield a decrease in decoding accuracy as a function of object distance (*H* = 3.53, *p* = 0.4735, Kruskal-Wallis test, *n* = 17 experiments). **j,** Same data as in the previous panel, but each colour shows a different experimental session.

**Supplementary Figure 7.**
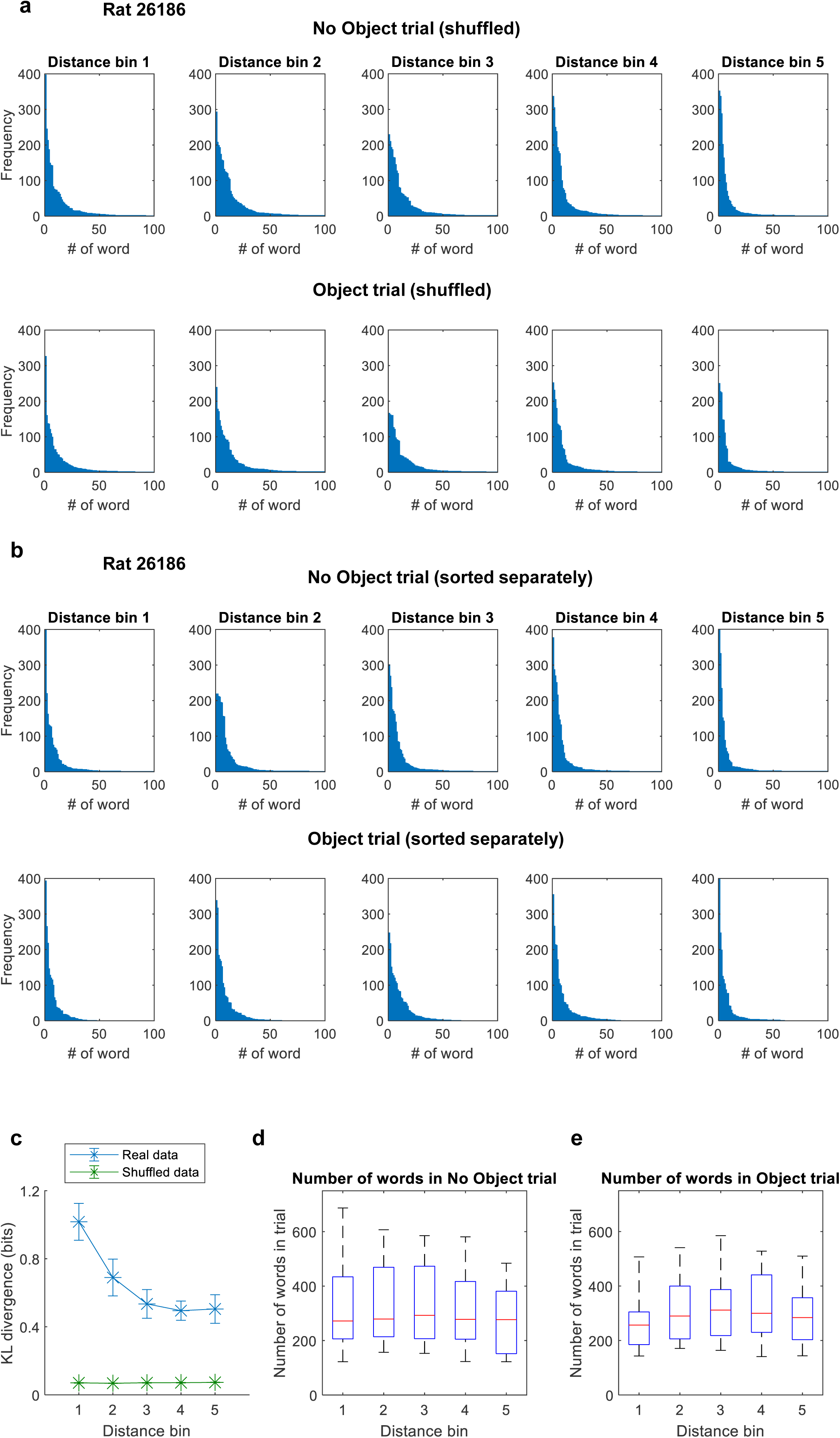
Controls related to word distributions. **a,** Example of word frequencies in shuffled data. Shuffling was performed by randomly assigning trial labels (No Object and Object) to data from No Object and Object trials. As in Figure 5c, distributions were sorted according to frequencies in the No Object trial (so word #1 at the top is the same as word #1 at the bottom). The bottom distribution has no “spikes”, suggesting that spikes in Figure 5d reflect reorganization of words in the actual data. **b,** Example of word frequencies from No Object (top) and Object (bottom) when these have been sorted independently according to frequencies within the same trial (No Object distributions sorted according to frequencies in No Object trial; Object distributions sorted according to frequencies in Object trial). Because of the independent sorting, word identities do not match between top and bottom (unlike in Fig. 5d). Shapes of distributions are nearly identical between top and bottom. Quantification for this panel is in Figure 5e. **c,** KL divergence between word distributions in No Object and Object trials as a function of distance of the animal relative to the object in real and shuffled data. The KL divergence is calculated between the top and bottom distribution for each distance bin. In shuffled data, the KL divergence is small and flat (green curve), meaning that word frequencies are nearly identical. The curve for real data is reproduced from Figure 5e for comparison (blue curve). **d,** Number of words used in each experiment in the No Object trial (n = 17 experiments). **e,** Number of words used in each experiment in the Object trial (n = 17 experiments). The number of words was the same across distance bins in both the No Object trial (*H* = 1.58, *p* = 0.8117, Kruskal-Wallis test) and Object trial (*H* = 2.62, *p* = 0.6227, Kruskal-Wallis test).

## STAR METHODS

### KEY RESOURCES TABLE

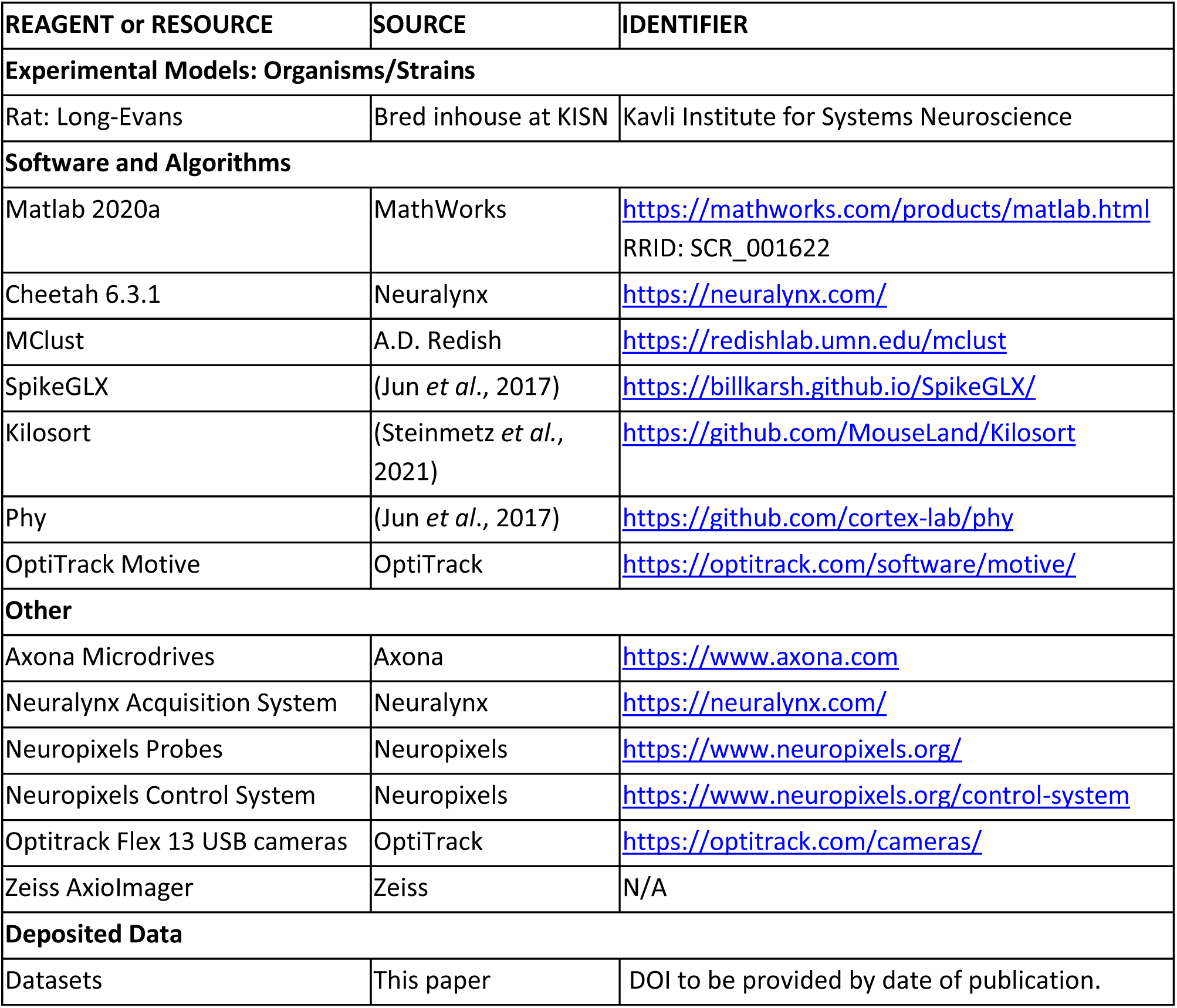

### RESOURCES AVAILABILITY

Further information and requests for resources should be directed to and will be fulfilled by the lead contact, May-Britt Moser.

#### Materials availability

This study did not generate new unique reagents.

#### Data and code availability

Datasets and custom codes used in this paper will be available at a publicly accessible repository by the date of publication (DOIs to be provided).

### EXPERIMENTAL MODEL AND SUBJECT DETAILS

Data were obtained from 10 male Long-Evans rats aged 2-3.5 months (320-470 grams) at the time of implantation. The rats were housed with 2-4 littermates prior to surgery. After implantation, the rats were housed individually on a single floor in large, dual-floor metal cages (95 x 63 x 61 cm) with access to food and water *ad libitum* throughout the entire experiment. The cages contained nesting material and were enriched with cardboard and plastic houses and contained a heightened platform. The rats were kept in a temperature and humidity controlled environment and kept on a 12 hour light/12 hour dark schedule. All testing occurred during the dark phase. All experiments were performed in accordance with the Norwegian Animal Welfare Act and the European Convention for the Protection of Vertebrate Animals used for Experimental and Other Scientific Purposes and were approved by the Norwegian Food Safety Authority (permit numbers 7163 and 18011).

### METHOD DETAILS

#### Surgery and electrode implantation

##### Tetrode experiments

Seven rats were implanted with one or two ‘microdrives’, each containing a single bundle of four or eight tetrodes. Five of these rats were implanted with a single eight-tetrode microdrive targeting the CA1 and CA3 areas of the hippocampus, whereas two rats were implanted with one four-tetrode microdrive targeting CA1/CA3 in one hemisphere and one eight-tetrode microdrive targeting the medial entorhinal cortex (MEC) of the opposite hemisphere. Few cells were recorded on the MEC tetrodes and the data from these tetrodes was not used. Choice of hemisphere varied across rats. Tetrodes were constructed from four twisted 17-μm polyimide-coated platinum-iridium (90–10%) wires (California Fine Wire). The electrode tips were plated with platinum to reduce electrode impedances to between 120 and 220 kΩ at 1 kHz.

Anaesthesia was induced by placing the rat in a closed glass induction chamber filled with 5 % isoflurane (air flow: 0.4 l/min; oxygen flow: 0.4 l/min). The rat was given subcutaneous injections of buprenorphine (Temgesic, 0.03 mg/kg), meloxicam (Metacam, 1 mg/kg), and atropine (0.05 mg/kg^−^). The local anaesthetic bupivacaine (Marcain, 1 mg/kg) was injected subcutaneously before the incision was made. After injections the rat was moved to a raised platform with a mask and a stereotactic frame and the head was secured in place with a set of ear bars. The rat’s body was resting on a heating pad (38°C) to ensure that the body temperature was maintained throughout the surgery. Isoflurane levels were gradually reduced to 0.5-1.5 % depending on the physiological condition of the rat, which was evaluated by reflex responses and breathing patterns.

After the incision had been made, the periost was removed and the craniotomies were drilled. Tetrodes targeting CA1/CA3 were implanted without an angle at 3.2 mm lateral to the midline and 3.8-4 mm posterior to bregma. Tetrodes targeting MEC were implanted at 4.5 mm lateral to the midline and 0.1 mm anterior to the transverse sinus edge, with an anterior angle of 25° relative to the bregma/lambda horizontal reference. The initial depth of the tetrode tips varied between 1500 and 2000 µm relative to the dura mater. The microdrives were secured to the skull using an adhesive (OptiBond, Kerr), dental cement (Meliodent), and 2-5 Jeweller screws (M1.4) that were placed in vertically drilled holes in the parietal bones. One or two screws were placed in the occipital bone over the cerebellum and were connected to the drive grounds.

After the surgery, the rat was left in a heated chamber (30 °C) for 30-60 minutes for the immediate recovery phase, after which it was transferred back to the home cage. Additional doses of buprenorphine were administered 8-12 and 24 hours after the first injection. Meloxicam was administered once every 24 hours for as long as was assessed necessary (usually 3-4 days). The rat was left to recover for 3-5 days before the experiments began.

##### Neuropixels experiments

Three rats were implanted bilaterally with four-shank Neuropixels 2.0 silicone probes^47,48^ targeting the CA1/CA3 in one hemisphere, and the MEC in the opposite hemisphere. The MEC data were not used in the present study due to the small number of cells and experimental sessions. Two rats were implanted with one probe in each hemisphere, whereas one rat was implanted with two probes in the left hippocampus and one probe in the right MEC. In this case, the two hippocampal-targeting probes were glued together at the probe base prior to surgery and the in total eight probe shanks were implanted in the same craniotomy. The hippocampal probe tips were lowered to 7000-7500 µm relative to the dura mater. The stereotactic coordinates and the surgical procedure were the same as described above for tetrode experiments and have been described in detail elsewhere^32^. The probes were secured to the skull using Optibond and Venus (Kulzer) and protected by fitting a modified falcon tube. After surgery and the initial recovery in the heating chamber, the rat was kept in the home cage for approximately 3 hours before experiments began. Post-operative analgesia was administered as described above. The MEC data were not used because of the small numbers of recorded cells.

#### Behavioural procedures

Before surgery, the rats were habituated to being handled by the experimenter and familiarised with the experimental task and environment during training sessions. Each rat underwent 7-15 training sessions, lasting at least 20 minutes each on separate days. During each training session, the rat foraged for randomly scattered corn puff crumbs in a 150 cm x 150 cm square arena with 50 cm tall black walls. The floor mat was either matt black or dark blue/green. Tetrode rats were not familiarised with the object prior to surgery, whereas Neuropixels rats had six or seven extra training sessions before surgery where the object was placed into the arena in order to avoid novelty effects of the object during recordings.

Tetrode sessions were performed in one of two different rooms. In one room, the arena was encircled by a set of lightproof blue curtains. A large white paper sheet was attached to the curtain on one side as a distal cue. The arena in the second room had curtains hanging on only one side of the arena and various distal cues were visible to the rat on the remaining sides. Neuropixels sessions were performed in a single room, where the arena was surrounded by a set of lightproof blue curtains on three sides. On the fourth side, the curtains were open towards a white wall on which a shelf containing the recording apparatus was placed.

During experimental testing, each recording session consisted of either three or four trials. In all trials, the rat was freely foraging for randomly scattered corn puff crumbs. Each trial typically lasted 20-30 minutes, depending on how long it took the rat to fully cover the extent of the arena. The first trial was always an empty arena (No Object). This was followed by a trial in which a colourful Duplo tower (52 cm x 10 cm x 10 cm) was placed at or close to the middle of the arena (Object). In the third trial, the Duplo tower was moved to another, semi-random location in the arena, usually towards one of the four corners (Object Moved; approximate range of distance from the corner: 35-55 cm). A fourth trial in which the object was removed was used to check for carry-over (hysteresis) of firing at the former object location (No Object’). In the few cases where the rat would stop exploring the arena, the fourth trial was omitted from the experiment (this happened to 8.5% of the cells). Between each trial, the rat was placed on a towel in a flowerpot on a pedestal next to the arena for 1-3 minutes and given a cookie with vanilla cream and access to a water bottle.

It has previously been shown that the activity of CA cells can change dynamically over time without changes in context^22–24^. To control for this, a separate set of experiments was performed, where cells were recorded as the rat foraged freely in the same open field arena during four consecutive trials without an object (Control sessions; Fig. 2b). All other parameters were the same as for sessions with the object.

In the tetrode experiments, recordings typically started 3-5 days after implantation. The tetrodes were turned down in steps of 50 µm per day. To determine if the tetrodes were in the CA1 pyramidal cell layer, cells were recorded as the rat explored the empty open field arena and the presence of theta modulation together with the presence of at least 5-10 place cells was used as criteria. When the criteria were met for the first time, the full experimental protocol was started. The tetrodes were subsequently moved down in steps of 50 µm per day at the most, and the experiment was repeated several times over the course of 1-2 months (up to 25 recording sessions per animal). In two of the rats, the tetrodes were subsequently lowered to the CA3 and experiments were repeated as for the CA1 recordings.

The Neuropixels data were obtained from a total of four recording sessions from the three rats. Two of the rats were only recorded once, whereas the third was recorded two times (first in CA1 and then in CA3) with a 3-day interval between the two recording sessions. All recording sessions were performed within 72 hours after recovery from surgery.

#### Recording procedures

##### Tetrode experiments

During electrophysiological recordings, the microdrives (with tetrodes) were connected to a Neuralynx data acquisition system (Neuralynx, Tucson, AZ; Neuralynx Digital Lynx SX) through a multichannel, impedance matching unity gain head stage. The output of the head stage was conducted via a lightweight multi wire tether cable and a slip-ring commutator. Unit activity was amplified by a factor of 3000-5000 and band-pass filtered from 600 to 6000 Hz. Spike waveforms above a threshold of 30 or 40 µV were timestamped and digitized at 32 kHz for 1 ms. Local field potential (LFP) signals were recorded from each tetrode, amplified by a factor of 250-1000, low-pass filtered at 300-475 Hz and sampled at 1800-2500 Hz. An overhead camera recorded the position of two light-emitting diodes (LEDs) on the head stage in order to track the rat’s movements at a sampling rate of 25 Hz. The diodes were separated by 4 cm.

##### Neuropixels experiments

Signals were recorded using a Neuropixels acquisition system as described previously^32,47,48^. Briefly, the spike band signal was amplified at a gain of 80, low-pass-filtered at 0.5 Hz, high-pass-filtered at 10 KHz, and digitalised at 30 kHz on the probe circuit board. The signal was then multiplexed and transmitted to a Neuropixels PXIe acquisition module via a 5 m tether cable made using twisted pair wiring. A 3D motion capture system consisting of six OptiTrack Flex 13 cameras and Motive software was used to track the rat’s movement at a sampling rate of ∼120 Hz. Five reflective markers glued to a rigid body was attached to the rat’s implant during recording. Randomised sequences of digital pulses were generated by an Arduino microcontroller. In order to synchronise the timestamps of the two recording systems, the pulses were sent to the Neuropixels acquisition system as direct TTL input and to the OptiTrack system via infrared LEDs that were placed on the edge of the arena.

#### Histology and reconstruction of electrode placement

The rat was placed in a closed glass induction chamber filled with 5% isoflurane (air flow 1.2 l/min) to induce anaesthesia. It was then given an overdose of Equithesin injected intraperitoneally and perfused intracardially with saline followed by 4% formaldehyde. The brain was removed and stored in formaldehyde. Frozen 30 µm sagittal sections were cut, mounted on glass, and stained with Cresyl violet (Nissl).

The final position of the tip of each tetrode or the placement of the probe shank was identified on photomicrographs obtained with an Axio Scan.Z1 microscope and ZEN software (Zen 2.6; Carl Zeiss). The tip of the tetrode was used to determine the anatomical location of the recorded cell. For the Neuropixels experiments, the depths of the probe shank tips were measured in ZEN software. The known position of active recording sites relative to the tip was then used to determine the location of the recorded cells. CA1 and CA3 recording locations were distinguished based on depth and activity profiles along the probe shanks. Recorded cells that were judged to be in the cortex overlying the CA1 were excluded from all analyses.

#### Spike sorting and cell-inclusion criteria

Spike sorting of tetrode data was performed offline using the graphical cluster-cutting software MClust (A.D. Redish, https://redishlab.umn.edu/mclust). Spikes were clustered manually in 2D projections of the multidimensional parameter space (consisting of waveform amplitudes and waveform energies), using autocorrelation and crosscorrelation functions as additional separation tools and separation criteria. To follow cells across recording sessions, clusters of successive sessions were compared and identified to be the same unit if the locations of the spike clusters in the multidimensional parameter space were stable. Spike sorting of Neuropixels data was performed using a version of KiloSort 2.5^48^ with customisations as described previously^32^.

For both types of experiments, all trials in a session were clustered together. Single units were discarded if more than 1% of their inter-spike intervals were less than 2 ms. Further, cells were excluded if they had a mean spike rate of less than 0.05 Hz or greater than 10 Hz across the full recording session.

### QUANTIFICATION AND STATISTICAL ANALYSIS

Data analyses were performed with custom-written scripts in MATLAB (Version 2020a). The study did not involve any experimental subject groups; therefore, random allocation and experimenter blinding did not apply and were not performed.

#### Firing rate maps and identification of firing fields

2D rate maps that displayed the firing rate as a function of the position of the animal in the arena were calculated for each trial separately. First, position estimates were binned into a square grid of 2.5 cm x 2.5 cm bins. Next, the firing rate in each position bin was calculated as the number of spikes recorded in the bin, divided by the time the rat spent in the bin. The 2D rate map was smoothed with a Gaussian kernel with standard deviation of 2 bins in both the *x* and the *y* direction. Only time epochs in which the animal was moving at a speed above 2.5 cm/s were used for constructing the rate maps and for spatial analyses.

Firing fields in the rate map were detected by iteratively applying the Matlab ‘contour’ function (Mathworks), starting from the cell’s peak firing rate until it reached 3 × median absolute deviation (MAD) of the firing rates of all bins in the rate map. Firing fields were defined as contiguous areas within a contour with a size of at least 225 cm^2^ and with a minimum firing rate of at least 1 Hz.

#### Spatial information content

To calculate the spatial information content in each cell’s firing, the rate map was used. First, the spatial information rate^49^ was computed as

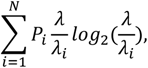

where N is the total number of bins (3200 bins in total), λ*_i_* is the firing rate in the *i*-th bin of the rate map, λ is the mean firing rate in the trial, and *P_i_* is the probability of the rat being in the *i*-th bin, calculated as the occupancy in the *i*-th bin (time spent in the bin) divided by the total duration of the trial. Spatial information content (in bits per spike) was then obtained for each trial by dividing the spatial information rate by the mean firing rate of the cell in that trial.

#### Shuffling of spike data

To determine significance for spatial information content, a per-trial shuffling procedure was implemented for each cell separately. For each trial, the sequence of spike timestamps was circularly shifted (with the end of the trial wrapped to the beginning of the trial) by a displacement that took random values between 20 seconds and 20 seconds less than the duration of the trial. Time shifts varied randomly between trials and between cells. The shuffling procedure was repeated 1000 times per trial per cell. For each shuffle iteration, the rate maps were recalculated and the spatial information content was computed using the position of the rat, which remained unchanged, and a shuffled distribution of spatial information content values was built. A cell was judged to be spatially modulated in the trial if the spatial information content calculated on the experimental data was above the 99^th^ percentile of the shuffled distribution.

#### Object score

To identify cells that fire at the location of the object, ‘object cells’, we calculated an ‘object score’ based on a template-matching procedure. First, we placed a Gaussian firing field at the object’s location, as a template for an ideal object cell. The standard deviation of the Gaussian firing field in the template was 25 cm, but we found that the results were similar for any reasonable standard deviation (field size). Then, we calculated the Pearson correlation between this template and the actual firing rate map of the cell. This was done separately both for the Object and Object Moved trials. Because an object cell should have high correlations with the template in both trials (i.e., the cell should consistently fire at the location of the object), we calculated the minimum value of the two Pearson correlations:

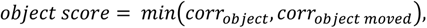

where *corr_object_* is the Pearson correlation between the template and actual firing rate map in the Object trial.

To obtain shuffled data, we chose the template’s location randomly, so that the template’s location could be anywhere in the arena (rather than at the object). After doing this in both the Object and Object Moved trials, we re-calculated the object score (500 times per cell). If the cell’s actual object score was larger than the 99^th^ percentile of the shuffled distribution, the cell was classified as an object cell.

#### Object-vector score and object-vector cell criteria

To identify cells firing at fixed distances and directions from objects, ‘object-vector cells’, we constructed ‘vector maps’ expressing each cell’s firing rate as a function of the animal’s distance and direction from the object^11,12^. We calculated the Pearson correlation between the two vector maps from the Object and Object Moved trials, and used the correlation as an ‘object-vector score’. Object-vector cells were defined as cells with an object-vector score exceededing chance levels determined by repeated shuffling of the experimental data. The shuffling was performed using the the spike-shifting procedure described above (‘Shuffling of spike data’) and implemented with 500 shuffle iterations for each of the object trials (Object and Object Moved). For each shuffle iteration, new vector maps were constructed and correlated with each other, resulting in a distribution of 500 shuffled correlation values per cell. A cell was judged to have an object-vector response if the vector-map correlation calculated on the experimental data was above the 99^th^ percentile of the shuffled distribution.

A cell was excluded from the analysis of object-vector cells (Fig. 1) if its potential object-vector field fell outside the arena in the Object Moved trial (e.g. if object was moved to the south-east corner, an object-vector cell firing at a south-east angle relative to the object would have fired outside the arena). This was assessed by identifying, for each cell, the centre-of-mass (COM) of all firing fields in the Object trial and the vectors between the COMs and the object. The vectors were then transferred to the Object Moved trial. If all COMs fell outside of the arena in the Object Moved trial, the cell was excluded from the analysis of object-vector cells. This amounted to 28.8% of all cells in CA1 (*n* = 342 cells). The remaining cells (*n* = 844 cells) were classified as object-vector cells if the following criteria were met:

1. The spatial information content was above the 99^th^ percentile of the cell’s shuffled distribution of spatial information content values in both the Object and the Object Moved trials. In the CA1 data, 78.2% of the cells met this criterion.
2. The object-vector score was above the 99^th^ percentile of the shuffled distribution for the cell. Of the 844 cells in the analysis, 10.5% (*n* = 89) had an object-vector score above the 99^th^ percentile.
3. At least one firing field was present in the vector maps from both the Object and the Object Moved trials. This was true for 91% of the cells that passed criteria 1 and 2 (*n* = 81 cells).
4. Finally, to distinguish object cells from object-vector cells, the COM of the object field was required to be at least 15 cm away from the centre of the object. 21 cells that passed criteria 1-3 were excluded because the firing field was closer than 15 cm from the object centre. After applying this criterion, 60 cells were accepted as object-vector cells (in total 7.1% of CA1 cells).

Unlike in previous publications on object-vector cells in the MEC^11,12^, cells were not excluded on the basis of having a firing field in the No Object trial.

#### Region-of-interest analysis for object-vector responses

To further confirm the difference between object-vector cells and other cells, an additional procedure for comparing object-vector fields across trials was implemented (Supplementary Fig. 3b, c). For object-vector cells with more than one field, the object-vector field was defined as the field with the highest similarity in distance and angle to the object between the Object and Object Moved trials. First, for each cell, firing fields in the Object trial were identified using the algorithm described above. A region-of-interest (ROI) was then defined as a circle with centre at the centre-of-mass of the field and radius calculated as

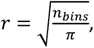

where n_bins_ is the number of bins in the rate map occupied by the field. The vector from the centre of the ROI to the centre of the object was calculated. The ROI and the vector were then transferred to the rate map from the Object Moved trial, using the object position of the Object Moved trial as reference (Supplementary Fig. 3b). To assess the similarity of firing within the ROIs in the Object and Object Moved trials, the firing rate overlap was calculated as the mean firing rate within the less active ROI divided by the firing rate in the more active ROI. To compare the ROI firing rate to the rest of the rate map, a shuffling procedure where the ROI was placed at random locations in the Object Moved rate map (‘Shuffled ROI’) was then applied and repeated 1000 times. Bins of the Shuffled ROI that fell outside the boundaries of the rate map were wrapped around to the opposite site of the rate map. For each iteration, the rate overlap between the actual ROI in the Object trial and the Shuffled ROI in the Object Moved trial was calculated. The mean across shuffle iterations was then calculated and used as the Shuffled ROI rate overlap for each cell (Supplementary Fig. 3c).

#### Spatial correlation and rate overlap

In order to determine whether the firing activity of single cells changed or remained stable when the object was introduced, moved, or removed from the environment, two measures were considered:

1. Spatial correlation
2. Rate overlap

The spatial correlation between two trials was defined as the Pearson correlation between the overlapping parts of the two firing rate maps (excluding bins that contained no value in either rate map). The rate overlap between two trials was calculated by dividing the mean firing rate in the less active trial by the mean firing rate in the more active trial.

To classify a cell’s response as changing or stable, a baseline stability of the cell’s firing within a trial (’within-trial’) was first assessed. The position data and the corresponding spike timestamps from the first trial (No Object) were binned into *n* sections of 1-second duration, where *n* is the full duration of the No Object trial in seconds. The bins were randomly assigned to two groups of size *n*/2. New rate maps for the two groups were then constructed, and the spatial correlation and the firing rate overlap between them was calculated. The procedure was repeated 500 times for each cell, with different bins randomly assigned to the two groups for each permutation.

For each between-trial comparison (e.g., No Object – Object), the same procedure as for the within-trial comparison was used. Position data and corresponding spiking data were randomly assigned to two groups of *n*/2 bins of 1 second duration for each of the two trials separately. Only one of the two groups (i.e., half of the data) from each trial was used and spatial correlation and rate overlap between the two trials were calculated as described above. The procedure was again repeated 500 times per between-trial comparison. A cell’s response was classified as changing if the mean of the between-trial distribution fell below the 1^st^ percentile of the within-trial distribution from the No Object trial, and if the mean ranks of the two distributions were judged to significantly different (assessed using a Wilcoxon signed-rank test).

#### Region-of-interest analysis for object traces

In order to investigate the presence of potential object-trace fields, firing fields were first identified in the rate map for No Object’. Then, a circular ROI was defined in the same way as for the object field analysis described in section ‘Region-of-interest analysis for object responses’. The ROI was then transferred to the same location in the rate maps for the other three trials (No Object, Object, and Object Moved; note that, unlike in the ROI analysis for object-vector responses, the object position was not used as a reference point). The firing rate overlap was calculated between the No Object’ ROI and the other three ROIs.

To determine whether an object trace was present, a shuffling procedure where the ROI identified in No Object’ was placed at random locations in the Object Moved rate map (‘Shuffled ROI’) was applied. Bins of the Shuffled ROI that fell outside the boundaries of the rate map were wrapped around to the opposite site of the rate map. This was repeated 1000 times, and for each shuffle iteration, the rate overlap between the actual ROI in the No Object’ trial and a Shuffled ROI in the Object Moved trial was calculated. All 1000 values of rate overlap were then used to build a shuffled distribution. If the rate overlap between the ROI in No Object’ and Object Moved was greater than the 99^th^ percentile of the shuffled distribution, the cell was classified as having an object-trace field (Supplementary Fig. 5).

#### Bayesian decoding algorithm

For this and all other sections using population activity, we used n = 15 tetrode sessions with a minimum of 25 and a maximum of 39 cells recorded simultaneously (the threshold for including a tetrode session being that it had a minimum of 25 cells) and n = 2 Neuropixels sessions with 620 cells (from two probes) and 135 cells (from a single probe), respectively.

To assess whether population activity in the hippocampus contains information about objects in the environment, we used a decoding approach. Building upon an existing Bayesian decoder^26^, we used rate maps to infer whether samples of neural activity occurred in the absence or presence of an object (No Object or Object trial). We divided the data into two groups, for training and testing (first half of data used for training and second half for testing; then vice versa in a separate iteration). While the training data was used to create rate maps, the testing data consisted of samples that the decoder classified as coming from the “No Object” or “Object” trial. Specifically, one sample of the testing data was a vector of spike counts from all hippocampal cells in a 1 second time window,

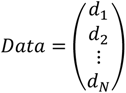

where *d_i_* is the number of spikes produced by each hippocampal cell in the 1 second time window and N is the total number of recorded cells in a session. Overall, our goal is to compare the likelihood that the “Data” are from the “No Object” trial against the likelihood that they are from the “Object” trial. This is given by the likelihood ratio:

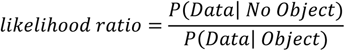

To calculate *P*(*Data*| *No Object*), because “Data” could have occurred when the animal was in any spatial bin, we started by introducing another term *x* on the right-hand side of the probability and asking how likely it is that the “Data” are from a specific spatial bin *x* (50 × 50 spatial bins, *x* = 1, 2…, 2500) in the No Object trial: *P*(*Data*| *x*, *No Object*).

To calculate the likelihood *P*(*Data*| *x*, *No Object*), we assume that each cell fires according to a Poisson distribution with mean firing rate *r_i_*(*x*), where *i* is the cell index and x is the spatial location of the animal. Each *r_i_*(*x*) is obtained by going to the rate map for cell *i* and extracting the firing rate in spatial bin *x*. In that sense, rate maps constructed from training data are used as a look-up table. The likelihood was calculated by,

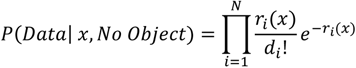

where *d_i_* is the number of spikes produced by cell *i* and where the multiplication over Poisson distributions from *N* cells indicates the assumption of independence between firing of different cells. Note that the more consistent the set of spike counts {*d_i_*} are with the set of firing rates {*r_i_*(*x*)}, the larger the value of the likelihood.

The likelihood was next multiplied by a prior,

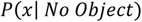

given by the fraction of time the animal spent in spatial bin *x* during the No Object trial. This accounts for the intuition that the “Data” are more likely to come from a given spatial bin if the animal spent a lot of time in that bin. We then marginalised over all spatial bins *x* (50 × 50 bins),

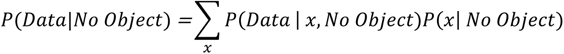

to obtain the overall likelihood of the No Object trial. The same calculation is performed for the Object trial to obtain the overall likelihood of the Object trial, *P*(*Data*| *Object*). The ratio of these terms determines the decoder’s output. That is, if 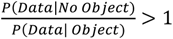, the decoder infers the “Data” as being from the No Object trial; otherwise, from the Object trial.

Since the length of an experimental trial was 1800 seconds, 1800/2 = 900 seconds could be used as testing data. With a bin size of 1 second, this gave us 900 testing samples. Given the two iterations described above (using either the first or second half of the trial as testing data), we had 900×2 = 1800 testing samples. The decoding accuracy was the fraction of 1800 samples classified correctly. The chance level of the decoder was calculated through a shuffling procedure (see “Shuffling trial labels”).

#### Calculation of decoding accuracy as a function of distance and allocentric angle

To understand how decoding performance depended on the animal’s spatial relationship with the object, we asked how decoding accuracy changed as a function of the animal’s distance and allocentric angle relative to the object (Fig. 4a-c and d-f, respectively). To examine the role of distance, we divided spike- and position data into 5 distance bins, forming concentric rings around the object (Fig. 4a). By creating training data and testing data for each specific distance bin and feeding these separately into the decoder, we could assess how decoding accuracy differed as a function of distance from object. For each experimental session, we optimized distance bins so that we had an equal number of testing samples within each bin. For example, if the animal spent little time near the object, distance bin 1 was made larger to obtain more samples. This helped to account for differences in the animal’s behavior in different experimental sessions. Specifically, the length of an experimental session was 1800 seconds, allowing 1800/2 = 900 seconds to be used for testing data. This gave 900/5 = 180 seconds of testing data per distance bin. Because the bin size was 1 second, we had 180 testing samples per distance bin. The decoding accuracy in each distance bin was the fraction of 180 testing samples classified correctly. To examine the role of allocentric angle, we used an analogous procedure. In this case, we divided spike- and position data into 5 angular bins, forming sectors of a circle (Fig. 4d), and created training and testing data separately for each angular bin.

#### Performance of decoder

##### Effect of bin size

To assess the performance of the decoder, we first asked how decoding performance depended on the bin size of testing data. For this analysis, we ran the decoder separately using bin sizes of 10, 50, 100, 500, 1000, 5000 and 10000 ms and calculated the decoding accuracy. Larger bin size corresponds to observing more spike counts from each hippocampal cell, which will lead to spike counts {*d_i_*} that are more similar to the mean firing rates {*r_i_*(*x*)}. Consistent with this idea, performance increased with increasing bin size. Specifically, performance increased rapidly up to 1000 ms, after which it saturated and gains of increasing bin size were lower (Fig. 3c). Based on these results, we chose 1000 ms as bin size for all main analyses.

##### Effect of number of cells

To assess how decoding performance depended on the number of cells fed into the decoder (Fig. 3e), we used a subsampling approach. We randomly selected a subset of *k* cells from a Neuropixels dataset with a total of N = 620 cells, iterating over *k* = {1, 5, 10, 20, 30, 50, 100, 200, 300, 400, 500} cells. For each *k*, we ran the decoder 100 times, using different subsets of cells each time.

#### Controls for results of decoder

##### Fixed distance bins

To ensure that our findings did not depend on using distance bins with an equal number of samples (described in “Calculation of decoding accuracy as a function of distance and allocentric angle”), we repeated the main analysis with fixed distance bins. In this case, we divided spike- and position data into distance bins with fixed values of 0-20 cm, 20-40 cm, 40-60 cm, 60-80 cm and 80-100 cm for all experimental sessions, ignoring differences in the animal’s behavior.

##### Decoding 1st half vs. 2nd half of No Object trial

To ensure that our findings could not arise without the presence of an object, we attempted to classify data samples belonging to either the 1st and 2nd half of the No Object trial, as a negative control. For this, we divided the No Object trial at its midpoint. For example, a 1800 seconds long trial would be split at 900 seconds, with 450 seconds for training data and 450 seconds for testing data for each one of the halves. The two halves were then used as input to the decoder (rather than the No Object and Object trials).

##### Shuffling trial labels

To ensure that our findings reflected a real relationship between population activity and trial type, we shuffled the trial labels of all testing data samples. That is, after shuffling, a spike count vector labelled “No Object” had 50% chance of being labelled “Object”. To perform the shuffling, we first concatenated testing data from the “No Object” and “Object” trials into an N×T spike matrix, where N is the number of cells and T the total number of timepoints. To shuffle the trial labels, we first randomly permuted the set of integers {1, 2, 3 … *T*} using the MATLAB function randperm. For example, consider the simple case of T = 10. The original arrangement of the N×1 spike count vectors in the matrix would be {1, 2, 3, 4, 5, 6, 7, 8, 9, 10}, where timepoints 1-5 would come from “No Object” and timepoints 6-10 would come from “Object”. The permuted rearrangement might be {7, 1, 2, 6, 10, 4, 8, 3, 9, 5}. After permutation, the first T/2 spike count vectors were labelled “No Object” and the last T/2 spike count vectors were labelled “Object”. This shuffling procedure destroys any real relationship between population activity and trial type.

##### Random partitioning into training and testing data

In the main analysis, we used the first half of the session as training data and the second half as testing data (and vice versa in a separate iteration). To confirm that our findings did not depend on such a partitioning, we used an alternative partitioning method where we randomly chose which set of timepoints would constitute training data. The random partitioning was analogous to the shuffling procedure above, but without concatenating spike matrices from the “No Object” and “Object” trials. For example, if data from “No Object” consisted of T timepoints, we generated a random permutation of the set of integers {1, 2, 3 … *T*} and used the first T/2 integers as timepoints for training data and the last T/2 integers for testing data (and vice versa in a separate iteration).

##### Decoding based on the animal’s occupancy

To ensure that our findings did not reflect differences in occupancy of animals across distance bins, we developed an alternative decoder that used the animal’s occupancy to try to infer the trial type of the animal. This decoder had a similar structure to the Bayesian decoding algorithm described above. However, each sample of testing data was now the instantaneous position of the animal in space,

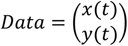

For each sample of testing data, the likelihood could be calculated by using the occupancy map as a look-up table. Specifically, we looked up which *x* and *y* bins in the occupancy map corresponded to the current sample of testing data. The probability that the testing sample came from the No Object trial was then proportional to the time spent by the animal inside those x and y bins,

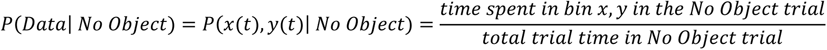

and similarly for the Object trial. The likelihood ratio could then be calculated as,

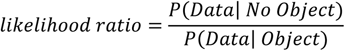

where we decoded the trial type as “No Object” if 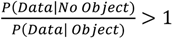 and “Object” otherwise.

#### Subsampling for decoder

To understand how widely distributed the object-centered population coding was across the recorded cells, we asked how many cells we needed to see the distance-based decrease in decoding accuracy (Fig. 6). To answer this, we used a subsampling procedure before feeding data into the decoder. We used Neuropixels data from a rat implanted with two probes in left hippocampus (N = 620 cells). The subsampling was analogous to the shuffling described under “Shuffling trial labels”. We took the set of all cells {1, 2, 3, …, *N*}, where each integer corresponds to a different cell identity, and randomly permuted the set using the MATLAB function randperm. We then took the first *k* integers of the permuted set, corresponding to *k* cells, and used them to create the subset. The number of subsampled cells, k, took the values 1, 5, 10, 30, 100, 200, 300, 400 and 500. For each k, we ran the decoder 20 times, (20 times being sufficient to get negligible error bars), so that the results were independent of cell identities and cell properties. In this way, we could determine how the number of cells influenced the decoding results (Fig. 5a) and show the distribution of distance coding across the entire hippocampal population.

#### Measuring strength of distance-specific reorganization

To estimate the strength of the distance-specific reorganization in populations of cells (and in individual neurons), we calculated two separate measures (Fig. 6d, f). Firstly, after having run the decoder and having obtained the curve showing decoding accuracy as a function of distance, we calculated

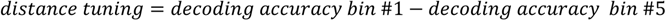

A strong distance-specific reorganization should be associated with large distance tuning. Secondly, we fitted a line to the same curve via least squares^50^,

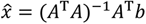

where *A* was the design matrix,

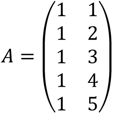

*b* was a vector containing the decoding accuracy of the cell in each distance bin *i*,

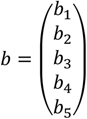

and 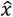 was a 2-element vector containing the intercept (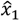) and slope 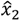 of the fitted line. A strong distance-specific organization should be associated with a more negative slope 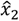. The results obtained using the two measures (distance tuning and slope) were consistent.

#### Population vector correlation

To assess the similarity between populations of rate maps from the No Object and Object trials, we calculated the correlation between stacks of rate maps from each trial. For each experimental session, we stacked the rate maps from the No Object trial and Object trial separately, while preserving the cell ordering. This procedure generated two N×m×n matrices, where N is the number of cells recorded in the session, m is the number of y-bins and n is the number of x-bins of the rate maps (*m* = *n* = 50 bins). We then calculated the Pearson correlation *r*(*x*, *y*) between two N-dimensional vectors of firing rates, one from each trial, for every spatial bin. After computing the Pearson correlations, we obtained an m×n map showing Pearson correlations as a function of space (Fig. 4g). After computing the correlation map, we allocated correlation values *r*(*x*, *y*) to distance bins and calculated the mean correlation in every distance bin (5 bins in total, bin size = 20 cm). Repeating this analysis for all experimental sessions (n = 17 from both tetrode and Neuropixels data), we were able to determine how correlations between the stacks changed as a function of distance from object (Fig. 4h).

#### Definition of words and counts

To understand which specific features of population activity explained the decrease in decoding accuracy as a function of distance from object, we considered two descriptions of population activity that we refer to as words and counts. These concepts have previously been used to study neural coding at the population level^27–29^. First, we expressed population activity in an N×T matrix, where N is the number of cells and T the number of timepoints in one trial. Each entry of the matrix corresponds to the number of spikes generated by each cell in specific time bin. To obtain mostly ones and zeroes (in rare cases, 2 or 3) we used 10 ms bins. We defined a specific pattern of activity and silence across the population as a “word”. We had as many words as timepoints sampled at 10 ms. The length of each word was the number of cells in the population (tetrode data: 25-39 cells; Neuropixels data: subsampled to contain 30 cells). In contrast, we defined the total number of spikes generated by the population in a time bin as the “count”. Words are a detailed description of population activity that preserves the cell identity, while counts are a coarse description of population activity. To illustrate, in a population of 4 neurons, the word [1, 0, 0, 1] would mean that the 1st and 4th neuron fired one spike in that time bin, while the 2nd and 3rd neurons remained silent. The total count would be 2 = 1 + 1. The word [0, 0, 0, 1] would mean that the 4th neuron fired one spike, while all others remained silent. The total count would be 1.

#### Mutual information

To quantify how much information words and counts provided about the presence of an object in the animal’s environment, we calculated the mutual information (MI)^51,52^ between the population response and the trial type (No Object and Object). The MI tells us how much information one variable (e.g., the words) provide about another variable (e.g., the trial type) independent of assuming any specific relationship between the two variables (e.g., the relationship could be completely non-linear). For words, the calculation was given by

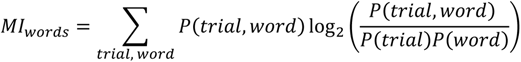

where *P*(*word*) and *P*(*trial*) are the marginal probabilities of the words and trial type and *P*(*trial*, *word*) is the joint probability. Similarly, for counts the calculations was given by

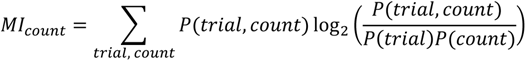

where *P*(*count*) and *P*(*trial*) are the marginal probabilities of the counts and trial type and *P*(*count*, *word*) is the joint probability. All probabilities were estimated from the frequency distributions. When calculating the MI of words, the number of bins was the same as the number of words in both trials (No Object and Object). For counts, the number of bins was determined by the maximum counts observed in both trials. For example, if the maximum number of spikes in a single time bin was 14, bins would range from 1 to 14 (in steps of 1). Being highly frequent and uninformative, the all-zero vector was excluded from our analyses and therefore the zero count was also excluded.

##### Bias-subtraction

Calculation of MI on datasets with small numbers of samples is well-known to lead to a bias where one overestimates the amount of information^30^. To ensure that our findings were not caused by sampling problems, we used a shuffling procedure to calculate the bias and subtract it from the mutual information. For this, we first shuffled trial labels as described for the decoder under “Shuffling trial labels”. After shuffling, a vector originally labelled “No Object” now had a 50% chance of being labelled “Object”. In other words, the shuffling destroys any relationship between the two variables (response and trial type) and shows the amount of “information” we should expect without such a relationship. The shuffling was performed 50 times, after which we calculated the mean MI across all 50 shuffles. This value (an estimate of the ‘bias’) was subtracted from the MI calculated on real data.

#### Word distributions

To directly visualize the change of words between No Object and Object, we calculated frequency distributions of words. For this, we first created a spike matrix based on activity from both trials (as in Fig. 5a, bin size = 10 ms, n = 25-39 cells). We then calculated the occurrences of each word, plotting the result in a histogram (Fig. 5d). To facilitate comparison between No Object and Object (for example, word #8 in No Object is the same as word #8 in Object), we applied the same sorting to the words in both trials. Specifically, we sorted words from No Object in order of descending frequency (Fig. 5d, top) and applied the same sorting to the Object trial (Fig. 5d, bottom).

To ensure that any differences in words between No Object and Object could not arise by chance, we randomly shuffled the trial labels (as described under “Shuffling trial labels”) and calculated word distributions on the shuffled data (Supplementary Fig. 7a).

Additionally, we wondered if the overall shapes of the word distributions changed between No Object and Object (for example, if more words are used in the Object trial, we would expect a longer tail in the distribution). We therefore sorted words from both trials in order of descending frequency (Supplementary Fig. 7b). In that case, word #8 in the No Object trial was not the same word as word #8 in the Object trial.

#### Kullback-Leibler divergence between word distributions

To quantify how strongly word distributions (as in Fig. 5d) differed between the No Object and Object trials, we calculated the Kullback-Leibler (KL) divergence^51,53^, which is given by,

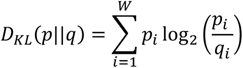

where *p_i_* is the probability of word *i* occurring in the No Object trial, *q_i_* is the probability of word *i* occurring in the Object trial, W is the total number of words from the No Object and Object trials, *p* = {*p_i_*} is the entire probability distribution of words from the No Object trial while *q* = {*q_i_*} is the entire probability distribution from the Object trial. Probabilities of the words were estimated in each trial as the normalized frequency distribution (number of times each word occurred/total number of timepoints). For visualization purposes, we plotted histograms of only the most 100 frequent words in each trial (Fig. 5d). However, the KL divergence was calculated using all words.

#### Statistical tests

All statistical analyses were performed in MATLAB (version 2020a). P-values are indicated in figures and legends. Wilcoxon signed-rank tests were used for paired data and Wilcoxon rank sum tests for unpaired data. Kruskal-Wallis tests were used for comparing differences between multiple groups. Binomial tests were used to compare fractions of observed cells (for example, object cells) to chance levels. Correlations were determined using Pearson’s product-moment correlation coefficients (*r*). The significance level was set to *p* = 0.05. All statistical tests were two-sided.

## REFERENCES

1. O’Keefe, J. & Nadel, L. The hippocampus as a cognitive map. (Clarendon Press; Oxford University Press, 1978).

2. O’Keefe, J. & Dostrovsky, J. The hippocampus as a spatial map. Preliminary evidence from unit activity in the freely-moving rat. (1971).

3. O’Keefe, J. Place units in the hippocampus of the freely moving rat. Exp Neurol 51, 78–109, doi:10.1016/0014-4886(76)90055-8 (1976).

4. Muller, R. U. & Kubie, J. L. The effects of changes in the environment on the spatial firing of hippocampal complex-spike cells. J Neurosci 7, 1951–1968 (1987).

5. Rivard, B., Li, Y., Lenck-Santini, P.-P., Poucet, B. & Muller, R. U. Representation of objects in space by two classes of hippocampal pyramidal cells. J Gen Physiol 124, 9–25, doi:10.1085/jgp.200409015 (2004).

6. Battaglia, F. P., Sutherland, G. R. & McNaughton, B. L. Local Sensory Cues and Place Cell Directionality: Additional Evidence of Prospective Coding in the Hippocampus. The Journal of Neuroscience 24, 4541, doi:10.1523/JNEUROSCI.4896-03.2004 (2004).

7. Lenck-Santini, P. P., Rivard, B., Muller, R. U. & Poucet, B. Study of CA1 Place Cell Activity and Exploratory Behavior Following Spatial and Nonspatial Changes in the Environment. Hippocampus 15, 356–369, doi:10.1002/hipo.20060 (2005).

8. Burke, S. N. et al. The influence of objects on place field expression and size in distal hippocampal CA1. Hippocampus 21, 783–801, doi:https://doi.org/10.1002/hipo.20929 (2011).

9. Hollup, S. A., Molden, S., Donnett, J. G., Moser, M.-B. & Moser, E. I. Accumulation of Hippocampal Place Fields at the Goal Location in an Annular Watermaze Task. The Journal of Neuroscience 21, 1635, doi:10.1523/JNEUROSCI.21-05-01635.2001 (2001).

10. Gothard, K. M., Skaggs, W. E., Moore, K. M. & McNaughton, B. L. Binding of hippocampal CA1 neural activity to multiple reference frames in a landmark-based navigation task. The Journal of Neuroscience 16, 823, doi:10.1523/JNEUROSCI.16-02-00823.1996 (1996).

11. Hoydal, O. A., Skytoen, E. R., Andersson, S. O., Moser, M. B. & Moser, E. I. Object-vector coding in the medial entorhinal cortex. Nature 568, 400–404, doi:10.1038/s41586-019-1077-7 (2019).

12. Andersson, S. O., Moser, E. I. & Moser, M.-B. Visual stimulus features that elicit activity in object-vector cells. Communications Biology 4, 1219, doi:10.1038/s42003-021-02727-5 (2021).

13. Deshmukh, S. S. & Knierim, J. J. Influence of local objects on hippocampal representations: Landmark vectors and memory. Hippocampus 23, 253–267, doi:10.1002/hipo.22101 (2013).

14. Geiller, T., Fattahi, M., Choi, J. S. & Royer, S. Place cells are more strongly tied to landmarks in deep than in superficial CA1. Nat Commun 8, 14531, doi:10.1038/ncomms14531 (2017).

15. GoodSmith, D. et al. Flexible encoding of objects and space in single cells of the dentate gyrus. Curr Biol, doi:10.1016/j.cub.2022.01.023 (2022).

16. Pastalkova, E., Itskov, V., Amarasingham, A. & Buzsáki, G. Internally generated cell assembly sequences in the rat hippocampus. Science 321, 1322–1327 (2008).

17. Meshulam, L., Gauthier, J. L., Brody, C. D., Tank, D. W. & Bialek, W. Collective Behavior of Place and Non-place Neurons in the Hippocampal Network. Neuron 96, 1178–1191.e1174, doi:https://doi.org/10.1016/j.neuron.2017.10.027 (2017).

18. Stefanini, F. et al. A Distributed Neural Code in the Dentate Gyrus and in CA1. Neuron 107, 703–716.e704, doi:https://doi.org/10.1016/j.neuron.2020.05.022 (2020).

19. Manns, J. R. & Eichenbaum, H. A cognitive map for object memory in the hippocampus. Learn Mem 16, 616–624, doi:10.1101/lm.1484509 (2009).

20. Leutgeb, S., Leutgeb, J. K., Treves, A., Moser, M. B. & Moser, E. I. Distinct ensemble codes in hippocampal areas CA3 and CA1. Science 305, 1295–1298, doi:10.1126/science.1100265 (2004).

21. Leutgeb, J. K. et al. Progressive transformation of hippocampal neuronal representations in “morphed” environments. Neuron 48, 345–358, doi:10.1016/j.neuron.2005.09.007 (2005).

22. Mankin, E. A. et al. Neuronal code for extended time in the hippocampus. Proc Natl Acad Sci U S A 109, 19462–19467, doi:10.1073/pnas.1214107109 (2012).

23. Mankin, E. A., Diehl, G. W., Sparks, F. T., Leutgeb, S. & Leutgeb, J. K. Hippocampal CA2 activity patterns change over time to a larger extent than between spatial contexts. Neuron 85, 190–201, doi:10.1016/j.neuron.2014.12.001 (2015).

24. Kentros, C. G., Agnihotri, N. T., Streater, S., Hawkins, R. D. & Kandel, E. R. Increased attention to spatial context increases both place field stability and spatial memory. Neuron 42, 283–295, doi:10.1016/s0896-6273(04)00192-8 (2004).

25. Vandrey, B., Duncan, S. & Ainge, J. A. Object and object-memory representations across the proximodistal axis of CA1. Hippocampus 31, 881–896, doi:10.1002/hipo.23331 (2021).

26. Posani, L. A.-O., Cocco, S., Ježek, K. & Monasson, R. Functional connectivity models for decoding of spatial representations from hippocampal CA1 recordings. (2017).

27. Schneidman, E., Berry, M. J., Segev, R. & Bialek, W. Weak pairwise correlations imply strongly correlated network states in a neural population. Nature 440, 1007–1012, doi:10.1038/nature04701 (2006).

28. Osborne, L. C., Palmer, S. E., Lisberger, S. G. & Bialek, W. The neural basis for combinatorial coding in a cortical population response. The Journal of neuroscience : the official journal of the Society for Neuroscience 28, 13522–13531, doi:10.1523/JNEUROSCI.4390-08.2008 (2008).

29. Schneidman, E. et al. Synergy from Silence in a Combinatorial Neural Code. The Journal of Neuroscience 31, 15732, doi:10.1523/JNEUROSCI.0301-09.2011 (2011).

30. Panzeri, S., Senatore, R., Montemurro, M. A. & Petersen, R. S. Correcting for the Sampling Bias Problem in Spike Train Information Measures. Journal of Neurophysiology 98, 1064–1072, doi:10.1152/jn.00559.2007 (2007).

31. Obenhaus Horst, A. et al. Functional network topography of the medial entorhinal cortex. Proceedings of the National Academy of Sciences 119, e2121655119, doi:10.1073/pnas.2121655119 (2022).

32. Gardner, R. J. et al. Toroidal topology of population activity in grid cells. Nature 602, 123–128, doi:10.1038/s41586-021-04268-7 (2022).

33. Fyhn, M., Hafting, T., Treves, A., Moser, M. B. & Moser, E. I. Hippocampal remapping and grid realignment in entorhinal cortex. Nature 446, 190–194, doi:10.1038/nature05601 (2007).

34. Yoon, K. et al. Specific evidence of low-dimensional continuous attractor dynamics in grid cells. Nat Neurosci 16, 1077–1084, doi:10.1038/nn.3450 (2013).

35. Alme, C. B. et al. Place cells in the hippocampus: eleven maps for eleven rooms. Proc Natl Acad Sci U S A 111, 18428–18435, doi:10.1073/pnas.1421056111 (2014).

36. McNaughton, B. L. & Morris, R. G. Hippocampal synaptic enhancement and information storage within a distributed memory system. Trends in Neurosciences 10, 408–415, doi:https://doi.org/10.1016/0166-2236(87)90011-7 (1987).

37. Leutgeb, S. et al. Independent codes for spatial and episodic memory in hippocampal neuronal ensembles. Science 309, 619–623, doi:10.1126/science.1114037 (2005).

38. Colgin, L. L., Moser, E. I. & Moser, M.-B. Understanding memory through hippocampal remapping. Trends in Neurosciences 31, 469–477, doi:https://doi.org/10.1016/j.tins.2008.06.008 (2008).

39. Latuske, P., Kornienko, O., Kohler, L. & Allen, K. Hippocampal Remapping and Its Entorhinal Origin. Frontiers in Behavioral Neuroscience 11, doi:10.3389/fnbeh.2017.00253 (2018).

40. Anderson, M. I. & Jeffery, K. J. Heterogeneous Modulation of Place Cell Firing by Changes in Context. The Journal of Neuroscience 23, 8827, doi:10.1523/JNEUROSCI.23-26-08827.2003 (2003).

41. Wills Tom, J., Lever, C., Cacucci, F., Burgess, N. & O’Keefe, J. Attractor Dynamics in the Hippocampal Representation of the Local Environment. Science 308, 873–876, doi:10.1126/science.1108905 (2005).

42. Jezek, K., Henriksen, E. J., Treves, A., Moser, E. I. & Moser, M.-B. Theta-paced flickering between place-cell maps in the hippocampus. Nature 478, 246–249, doi:10.1038/nature10439 (2011).

43. Kubie, J. L., Levy, E. R. J. & Fenton, A. A. Is hippocampal remapping the physiological basis for context? Hippocampus 30, 851–864, doi:https://doi.org/10.1002/hipo.23160 (2020).

44. Kelemen, E. & Fenton, A. A. Dynamic Grouping of Hippocampal Neural Activity During Cognitive Control of Two Spatial Frames. PLOS Biology 8, e1000403, doi:10.1371/journal.pbio.1000403 (2010).

45. Renaudineau, S., Poucet, B. & Save, E. Flexible use of proximal objects and distal cues by hippocampal place cells. Hippocampus 17, 381–395, doi:https://doi.org/10.1002/hipo.20277 (2007).

46. Yuste, R. From the neuron doctrine to neural networks. Nat Rev Neurosci 16, 487–497, doi:10.1038/nrn3962 (2015).

47. Jun, J. J. et al. Fully integrated silicon probes for high-density recording of neural activity. Nature 551, 232–236, doi:10.1038/nature24636 (2017).

48. Steinmetz, N. A. et al. Neuropixels 2.0: A miniaturized high-density probe for stable, long-term brain recordings. Science 372, doi:10.1126/science.abf4588 (2021).

49. Skaggs, W. E., McNaughton, B. L. & Gothard, K. M. in Advances in Neural Information Processing Systems 5 [NIPS Conference] 1030–1037 (Morgan Kaufmann Publishers Inc., 1993).

50. Agresti, A. Foundations of Linear and Generalized Linear Models. (John Wiley & Sons Inc, 2015).

51. Cover, T. M. & Thomas, J. A. Elements of Information Theory. (John Wiley & Sons, Inc, 2006).

52. Shannon, C. E. A Mathematical Theory of Communication. Bell System Technical Journal 27, 379–423, doi:https://doi.org/10.1002/j.1538-7305.1948.tb01338.x (1948).

53. Kullback, S. & Leibler, R. A. On Information and Sufficiency. The Annals of Mathematical Statistics 22, 79–86, doi:10.1214/aoms/1177729694 (1951).

